# Direction-sensitive neural correlates of voluntary pain regulation: regional response attenuation during down-regulation and exploratory prefrontal-valuation coupling

**DOI:** 10.64898/2026.10.01.756083

**Authors:** Meysam Kharghani, Elahe Gholami

## Abstract

Voluntary cognitive regulation can increase or decrease pain, but direction-sensitive neural correlates may emerge at different analytical levels. We reanalyzed 3,201 thermal-stimulation trials from 33 participants, integrating 0-200 warmth/pain ratings, regional fMRI activity, and trial-wise lagged coupling. Both instructions shifted ratings in the intended direction, with no reliable difference in mean success. Down-regulation was associated with lower bilateral S2/opercular and insular activity and higher left dlPFC/IFJ activity after FDR correction across 32 regions; run-matched comparisons were weaker and none survived the corresponding correction. Contrast-map Neurologic Pain Signature attenuation did not replicate in a rating-adjusted FWL-LSS trial-wise analysis despite strong temperature sensitivity. Up-regulation success covaried with an exploratory, post-selection lagged dlPFC/IFJ-valuation composite (r = -0.546, 95% CI [-0.764, -0.270]). The association exceeded matched reverse-direction, zero-lag, and time-reversed controls, lay outside permutation and circular-shift nulls, and persisted after AR(1) prewhitening. A transition-level analysis across 297 up-regulation transitions preserved a negative success-by-source interaction (adjusted three-target mean beta = -0.163, cluster-bootstrap 95% CI [-0.290, -0.054], Freedman-Lane p = 0.010). Six session-wise DCM variants showed no consistent regulation-versus-passive effective-connectivity effect and did not independently validate an up-specific directed mechanism. These analyses identify a down-regulation-associated regional pattern and a candidate prefrontal-valuation coupling association requiring independent confirmation.

## Introduction

Pain reflects interactions among nociceptive input, attention, expectation, affective meaning, and valuation (Atlas and Wager, 2012; Melzack and Wall, 1965; Roy et al., 2012; Tracey and Mantyh, 2007). Voluntary cognitive regulation provides a direct way to test these influences because comparable noxious stimulation can be intentionally amplified or attenuated. Treating regulation only as an up-versus-down contrast, however, obscures how each direction differs from passive experience and whether regulation is expressed in average regional responses, individual differences in interregional coupling, or both. Direction-resolved analyses are therefore useful without assuming that amplification and attenuation are mirror images or separate mechanisms.

The original analysis of OpenNeuro ds000140 used the Neurologic Pain Signature (NPS; Wager et al., 2013) to separate nociceptive from evaluative contributions to pain. Stimulus-intensity effects were mediated by NPS, whereas self-regulation was linked to a nucleus accumbens-ventromedial prefrontal cortex (vmPFC) pathway and did not reliably alter NPS expression in the canonical up-versus-down contrast (Woo et al., 2015). This left two questions relevant to pain modulation: whether either direction shows regional changes relative to passive stimulation, and whether individual regulatory success relates to coupling between cognitive-control and valuation systems.

Down-regulation may recruit lateral prefrontal regions involved in cognitive control and pain reappraisal (Adamczyk et al., 2020; Wiech et al., 2006) while attenuating secondary somatosensory/opercular and insular responses implicated in sensory-discriminative and nociceptive-affective processing (Petre et al., 2022; Tracey and Mantyh, 2007). Variation in up-regulation success may instead be expressed in coordination between dorsolateral prefrontal/inferior frontal junction (dlPFC/IFJ) control regions and vmPFC/medial orbitofrontal valuation regions. Pain-control and expectation studies support direction-sensitive temporal and regional effects, but agency, controllability, expectation, and voluntary regulation are related rather than interchangeable constructs (Büchel, 2023; Habermann and Büchel, 2025; Mourkojannis et al., 2025; Wittkamp et al., 2024).

Here, we quantified behavioral regulation in both directions, tested whether apparent baseline-state dependence was robust to alternative outcome definitions and change-score nulls, and evaluated direction-resolved regional fMRI effects. We then examined whether regulation success was associated with rating-adjusted trial-wise lagged prefrontal-valuation coupling, with matched temporal controls, AR(1) sensitivity, and a transition-level analysis. Whole-brain mapping, NPS analyses, and six run-separated dynamic causal models assessed the scope and convergence of the findings. Targeted regional analyses were not preregistered, and the coupling pathway was selected and characterized in the same sample; coupling and DCM results are therefore exploratory.

## Methods

### Dataset, participants and task

We reanalyzed the OpenNeuro dataset ds000140, version 00001 (Woo et al., 2018), hosted by the OpenNeuro data-sharing platform (Markiewicz et al., 2021) and distributed in Brain Imaging Data Structure format (Gorgolewski et al., 2016). The released sample comprised 33 healthy, right-handed participants (22 female, 11 male; age 19–55 years, M = 27.73, SD = 8.92).

The released demographic field encoded 22 participants as female and 11 as male. The source metadata did not document how sex was ascertained, and gender identity was unavailable. Because this secondary sample was not designed or powered for sex-stratified inference, sex was reported descriptively rather than included in the primary models.

Thermal stimulation was delivered to the volar surface of the left inner forearm using a TSA-II Neurosensory Analyzer (Medoc Ltd.) fitted with a 16-mm Peltier thermode end-plate; two sites on the middle forearm were alternated between runs (Woo et al., 2015). Each 12.5-s stimulus comprised a 3-s ramp-up, 7.5 s at the target temperature, and a 2-s ramp-down (Woo et al., 2015). Participants then provided an 11-s rating after each stimulation. The released event files contained temperature values ranging from 40.8 to 47.3, whereas the original report described six nominal stimulation levels from 44.3 to 49.3°C (Woo et al., 2015). Because the released metadata do not document a conversion between these scales, we analyzed the event-coded values as provided and did not treat them as directly equivalent to the nominal protocol temperatures. The combined warmth/pain rating variable (0–200) was likewise used without recoding; throughout this report, “rating” refers to this released composite rather than to a pain-only measure. Runs 1, 2, 4, 5, 6, 8 and 9 were passive-experience runs with 11 stimulation trials each. Runs 3 and 7 were regulation runs with 10 trials each, during which participants used imagery and reinterpretation to increase attention to the unpleasant and burning qualities of the stimulus or to decrease pain by focusing on pleasant warmth, cooling and numbing sensations (Woo et al., 2015). Regulation order was counterbalanced: 16 participants completed up-regulation first and 17 completed down-regulation first.

The behavioral dataset contained 3,201 trials: 2,541 passive trials, 330 up-regulation trials and 330 down-regulation trials. There were no missing subjective ratings or event-coded temperature values (Table 1).

**Table 1.** Sample, task and main imaging dataset.

| Measure | Value | Notes |
| --- | --- | --- |
| Participants | 33 | 22 female; 11 male; age 19–55 years |
| Behavioral trials | 3,201 | 2,541 passive; 330 up-regulation; 330 down-regulation |
| Functional runs acquired | 297 | Nine runs per participant |
| Functional runs retained for main imaging analyses | 288 | 222 passive; 33 up-regulation; 33 down-regulation |
| Functional runs excluded | 9 | All were passive runs and exceeded at least one severe motion criterion |
| Regulation runs retained | 66/66 | All 33 up-regulation and 33 down-regulation runs |

### Behavioral reconstruction and task validation

Each stimulation event was paired with the subsequent rating event from the same run. Temperature and rating values were analyzed on their released scales and after standardization within participant. Trial position was centered within run, run position was centered within participant, and a context indicator identified passive runs 5 and 6, which used a shifted temperature context in the original design (Woo et al., 2015).

Task validation used participant fixed-effects regressions with participant-clustered standard errors. The standardized-rating model included indicators for up- and down-regulation, within-participant standardized temperature, within-run trial position, participant-centered run position, the passive-context indicator and participant fixed effects. The corresponding raw-rating model used the same predictors. Finite-cluster sensitivity analyses used CR2 standard errors with Satterthwaite degrees of freedom and 9,999 null-imposed Rademacher wild-cluster-bootstrap replications.

### Passive expected pain and regulation success

Expected pain was estimated from passive trials using separate raw and standardized participant fixed-effects models. Each model included within-participant standardized temperature, within-run trial position, linear and quadratic participant-centered run terms, and the passive-context indicator. Direction-corrected trial-wise regulation success was defined as:

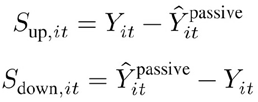

Thus, positive values indicated change in the instructed direction for both conditions. Participant mean success was assessed with two-sided one-sample t tests, sign-flip tests and bootstrap confidence intervals, and the up-minus-down comparison was computed from participant-level direction-corrected means.

### State-dependence and coupling-null analyses

The main state-dependence model regressed standardized regulation success on direction, passive-expected pain and their interaction, with temperature, trial position, run position, regulation order and participant fixed effects included as covariates. Sensitivity analyses varied the outcome definition or passive baseline, including raw success, observed ratings, bounded-logit ratings, available-room-normalized success, cross-fitted passive baselines, expected-pain and temperature bins, and boundary-trimmed samples.

Coupling-null simulations tested whether the observed direction-by-expected-pain association could arise from using expected pain in both the predictor and the change-score outcome. Ratings were generated from a model without a true direction-by-state interaction, success scores were recomputed and the interaction model was refitted. Two thousand simulations were performed for unbounded generated ratings and for ratings clipped to the observed 0–200 range. For each null distribution, we report its mean, median, central 95% interval and the percentile rank of the observed raw up-minus-down slope. Because the simulated null distributions were not centered at zero, absolute-value two-sided p values were not interpreted.

### fMRI preprocessing and quality control

Functional images were acquired on a 3-T Philips Achieva TX scanner using gradient-echo echo-planar imaging (TR = 2,000 ms, TE = 20 ms, flip angle = 72°, field of view = 224 mm, 64 × 64 matrix, 3-mm isotropic voxels, 42 interleaved slices, SENSE factor = 1.5) (Woo et al., 2015). The released functional data were preprocessed with fMRIPrep 25.0.0 using Docker (Esteban et al., 2019) and normalized to MNI152NLin2009cAsym space. Run-level quality control assessed the availability and dimensional consistency of preprocessed BOLD images, brain masks, and confound files, together with framewise displacement (FD) and confound summaries. Documented hard-exclusion criteria were missing preprocessed BOLD data, a missing brain mask, a mismatch between BOLD and confound lengths, mean FD greater than 0.50 mm, more than 30% of volumes with FD greater than 0.5 mm, more than 10% of volumes with FD greater than 1.0 mm, or maximum FD greater than 5.0 mm.

The dataset contained 297 functional runs. Nine passive runs exceeded at least one severe motion criterion and were excluded; no run was excluded for missing files or dimensional mismatch. The main imaging set therefore contained 288 runs (222 passive, 33 up-regulation and 33 down-regulation), and all 66 regulation runs were retained. Visual inspection of the retained preprocessed data indicated acceptable brain coverage and spatial alignment.

### First-level and group-level GLM

BOLD images retained for the univariate GLM were smoothed with a 6-mm full-width-at-half-maximum Gaussian kernel. First-level models were fitted with Nilearn 0.13.0 (Abraham et al., 2014) using an SPM hemodynamic response function and temporal derivative, cosine drift terms with a 180-s high-pass cutoff, an AR(1) noise model, a rating-period nuisance regressor and condition-specific within-run-centered temperature modulators when estimable. Nuisance regressors comprised the six rigid-body motion parameters, their temporal derivatives, squares of both sets (24 motion terms), white-matter and cerebrospinal-fluid signals, the first five available anatomical CompCor components, custom spike regressors for volumes with framewise displacement greater than 0.5 mm and available non-steady-state outlier regressors.

Run-level estimates were combined within participant by inverse-variance weighting:

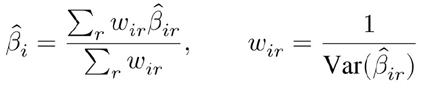

Group models were fitted with Nilearn SecondLevelModel and thresholded with voxel-wise false-discovery-rate correction at q < 0.05 (Benjamini and Hochberg, 1995). Motion robustness was evaluated with stricter run-level exclusions and exclusion of high-motion participants.

### ROI and multivariate-pattern analyses

Thirty-two targeted masks represented sensory and pain-related, control and valuation, subcortical and contextual regions. Cortical masks were constructed from the volumetric extended Human Connectome Project multimodal parcellation atlas (HCPex, 2-mm version) (Huang et al., 2022) and included bilateral S1, S2/operculum, whole, anterior and posterior insula, dACC/MCC, pgACC/rACC, vmPFC, medial orbitofrontal cortex, dlPFC/IFJ and SMA/pre-SMA. Subcortical masks were derived primarily from the Melbourne Subcortex Atlas, scale 4, 3-T, subcortex-only version (Tian et al., 2020); caudate masks used HCPex labels. The periaqueductal gray was defined as a 4-mm-radius sphere centered at MNI [0, −32, −10], and the exploratory hypothalamus mask combined probabilistic subunit maps (Billot et al., 2020). The full label-to-ROI mapping and voxel counts are provided in the repository.

Participant-level fixed-effect contrast estimates were averaged within each ROI and tested against zero using two-sided one-sample t tests across participants. Percentile bootstrap confidence intervals were based on 5,000 participant-level resamples. Ten ROI effect families were evaluated per imaging set: down-minus-passive, up-minus-passive, up-minus-down, regulation-mean-minus-passive, condition-specific stimulation estimates for passive, up, and down, and the corresponding three temperature-modulation estimates. Benjamini–Hochberg correction was applied across all 32 ROIs separately within each effect family; no correction was pooled across the ten families. The focal main-text regional result was the down-minus-passive family. For the run-position sensitivity analysis, Benjamini–Hochberg correction was likewise applied across all 32 ROI features separately within each regulation-direction-by-comparator family (two directions × four comparator definitions; eight 32-test families). The targeted ROI analyses were theory-guided but not preregistered and are therefore interpreted as non-confirmatory within-dataset analyses.

Expression of the published Neurologic Pain Signature (NPS) (Wager et al., 2013) was evaluated using two complementary paths. The contrast-map path calculated weighted-mean NPS expression from participant-level fixed-effect maps. The trial-wise path was re-estimated on 6-mm-smoothed, QC-retained BOLD data from all 288 runs using rating-adjusted FWL-LSS, yielding 3,102 stimulation-trial estimates. For each target trial, the nuisance design included a pooled regressor for all other stimulation events, a pooled rating-period regressor, the curated run-level confounds, and cosine drift terms with a 0.008-Hz high-pass cutoff; event regressors used the SPM hemodynamic response function. The target regressor and the NPS-projected BOLD time series were residualized against the nuisance design using an SVD-based pseudoinverse before estimating the target coefficient. Weighted-mean NPS expression was the primary scale; dot-product and L2-normalized forms were proportional rescalings and produced identical inferential statistics.

### Rating-adjusted trial-wise ROI estimation for coupling

Coupling-specific trial-wise regional estimates were obtained with a least-squares-separate (LSS) procedure and Frisch-Waugh-Lovell (FWL) residualization applied to unsmoothed, QC-retained BOLD data from all 66 regulation runs. The analysis yielded estimates for 660 stimulation trials in left dlPFC/IFJ, vmPFC, mOFC, and a combined vmPFC/mOFC mask. For each target trial, the nuisance design contained a pooled regressor for all remaining stimulation events, a pooled rating-period regressor, the 24-parameter motion expansion, white-matter and cerebrospinal-fluid signals, the first five available anatomical CompCor components, custom FD spike regressors, available non-steady-state outlier regressors, and cosine drift terms with a 0.008-Hz high-pass cutoff. Event regressors were convolved with the SPM hemodynamic response function; non-finite confound values were set to zero, and constant or all-zero columns were removed. For target-trial regressor x, nuisance design N, and ROI time series Y, the FWL estimate was:

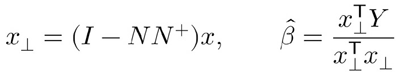

Here, N⁺ denotes the Moore–Penrose pseudoinverse of N. The SVD-based pseudoinverse used a relative cutoff of 10⁻¹², and the target regressor was required to retain unique residual variance after nuisance projection. Unlike the main first-level GLM, which used Nilearn run-wise mean signal scaling (signal_scaling = 0), this LSS path used unsmoothed fMRIPrep ROI time series in native intensity units without run-wise mean scaling. Consequently, run-matched LSS beta differences are not numerically comparable with the mean-scaled first-level GLM contrasts reported in the regional analysis. The primary coupling estimates did not impose temporal prewhitening. As a sensitivity analysis, all 660 trial estimates were re-estimated with the same event and nuisance specification using Nilearn run_glm with an AR(1) noise model (100 bins), and the complete matched coupling-control analysis was repeated. All 660 target regressors passed the predefined estimability criterion; design diagnostics and per-run coefficients are archived in the reproducibility materials.

### Lagged ROI-to-ROI coupling and pathway screening

Consecutive trial transitions were constructed only within runs. Each 10-trial regulation run contributed nine transitions, yielding 594 candidate within-run transitions in total (297 per regulation direction). For each participant and target, the standardized lagged model was:

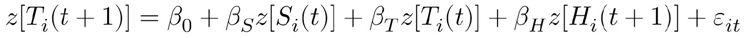

Here, S denotes the source ROI, T the target ROI, and H the event-coded heat temperature. Valid within-run transitions were pooled within participant and direction and estimated by ordinary least squares. With an intercept and three predictors, each participant-direction coefficient was estimated from nine observations and five residual degrees of freedom; individual coefficients may therefore be unstable. The source coefficient β_S is a temporally lagged statistical association and is not interpreted as causal or physiological directionality.

The exploratory targeted screen evaluated four theory-guided lagged pathway composites linking nucleus accumbens with vmPFC, anterior insula with dACC/MCC, left dlPFC/IFJ with valuation regions, and pgACC/rACC with periaqueductal gray. Each pathway quantity was correlated with 29 participant-level behavioral variables. Benjamini–Hochberg FDR was applied separately within each connectivity modality and pathway quantity; for up-regulation lagged coupling, this family comprised 116 correlations (four pathway composites × 29 behavioral variables). The archived materials did not establish a preregistered priority among the correlated success outcomes or a prespecified sequence for constructing the selected composite. Accordingly, the association between standardized up-regulation success and the left dlPFC/IFJ– valuation composite is treated as exploratory and post-selection.

The participant-level coupling score summarized the three related valuation-target definitions as an equal-weight composite:

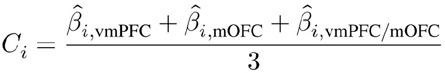

The vmPFC, mOFC, and combined vmPFC/mOFC masks are overlapping or nested and do not constitute independent anatomical replications. Post-selection characterization included Pearson and Spearman correlations; a covariate-adjusted standardized regression including regulation order, passive mean rating, and the participant-specific passive sensory slope, with HC3 standard errors; participant-label permutation; bootstrap confidence intervals; leave-one-participant-out estimates; and Cook-distance diagnostics. Matched controls used the same rating-adjusted trial estimates, within-participant standardization, and composite construction for reverse-direction, synchronous (zero-lag), and time-reversed models. A shared-source within-run permutation reused one source permutation per participant, run, and iteration across the three target definitions (5,000 iterations; seed 1234); a complementary nonzero circular shift preserved the within-run source sequence while changing its trial alignment (5,000 iterations; seed 8241). Participant bootstrap tests compared the forward correlation directly with each control (10,000 resamples; seed 2024), with FDR correction across the three comparisons. These follow-ups were conditional on the selected pathway and did not repeat or correct the screening stage.

To assess whether the selected association depended on noisy participant-specific coefficients estimated from only nine transitions, we performed a transition-level sensitivity analysis using the archived rating-adjusted AR(1)-prewhitened trial estimates. For each valuation target, Target(t+1), Source(t), Target(t), and Temperature(t+1) were standardized within participant across the nine up-regulation transitions. The unadjusted model fitted all 297 transitions directly as:

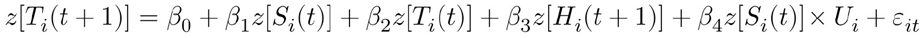

where U_i is standardized participant-level up-regulation success and β₄ is the success-by-source interaction of interest. Participant-clustered inference treated the participant as the independent cluster. The adjusted specification additionally included Source × regulation-order, Source × passive-mean-rating, and Source × passive-sensory-slope interactions. The equal-weight mean of the three target-specific β₄ coefficients was evaluated with 10,000 participant-label permutations, 10,000 participant-cluster bootstrap resamples, leave-one-participant-out analyses, and a Freedman-Lane residual-permutation test conditional on the three covariates. Random-slope mixed models were examined as diagnostics but were not used for primary inference when convergence or boundary warnings occurred. This sensitivity analysis was conditional on the selected pathway and did not repeat the 116-correlation screening stage.

### Session-wise dynamic causal modeling

Dynamic causal modeling and parametric empirical Bayes were performed in SPM25 (Wellcome Centre for Human Neuroimaging, London, UK) under MATLAB R2023b (The MathWorks, Inc., Natick, MA, USA) (Friston et al., 2003; Zeidman et al., 2019). Six exploratory five-node variants spanning prefrontal–valuation, affective–descending, and sensory–interoceptive architectures were estimated. Each retained functional run was modeled as an independent DCM, without concatenating time series or carrying neuronal and hemodynamic states across run boundaries. This produced 288 DCMs per variant (222 passive, 33 up-regulation, and 33 down-regulation) and 1,728 run-specific DCMs across the six-model family. Mean time series were extracted from run-specific or BOLD-resampled ROI masks using unsmoothed QC-retained data and standardized within run. Heat events were encoded at microtime resolution (16 bins per TR; reference bin 8), drove every selected node through the C matrix, and modulated each theory-permitted A-matrix edge through a single run-level B matrix. The nuisance matrix contained an intercept, a 128-s discrete-cosine high-pass basis, an HRF-convolved rating-period regressor, and all variable numeric columns from the curated run-level confound table after standardization and orthogonalization. The echo time was fixed at 20 ms.

Within each participant, a second-level PEB across all available run-specific DCMs estimated PassiveAtMeanRun, UpMinusPassive, DownMinusPassive, and linear RunPosition effects; a separate two-session PEB estimated DirectionMean and UpMinusDown. Third-level PEB with Bayesian model reduction and averaging was then applied to B-matrix parameters. Because direction-specific success scores and passive mean pain were strongly collinear when entered together, scientific interpretation used reduced, full-rank group designs that separated intercept-only condition effects, direction-specific success with regulation order, mean regulation success, and baseline/context covariates. Across these reduced designs, the maximum variance-inflation factor was 1.311. A parameter was described as retained only when its 90% posterior credible interval excluded zero and its BMA parameter-inclusion probability was at least 0.95. The six anatomical variants were treated as exploratory model-family sensitivities rather than a confirmatory multiple-comparison family; free energies were not compared across variants because the models used different regional time series.

### Statistical inference

Unless stated otherwise, tests were two-sided. Behavioral regressions used participant-clustered standard errors based on 33 participant clusters. Finite-cluster sensitivity analyses used CR2 standard errors with Satterthwaite degrees of freedom and 9,999 null-imposed Rademacher wild-cluster-bootstrap replications; intercept-only mean tests used the algebraically equivalent participant-cluster sign-flip implementation. Participant mean tests used t statistics with 32 degrees of freedom together with nonparametric or resampling checks where specified. ROI contrasts were assessed with two-sided one-sample t tests and 5,000-resample participant-level percentile bootstrap intervals. ROI and signature analyses used Benjamini-Hochberg false-discovery-rate correction within the explicitly defined correction families; correction was not pooled across the ten ROI effect families. Pathway screening used false-discovery-rate correction within modality-by-quantity families, and matched-control bootstrap comparisons used false-discovery-rate correction across the three direct contrasts (Benjamini and Hochberg, 1995). For DCM, posterior credible intervals and Bayesian-model-averaged parameter-inclusion probabilities were used as described above. Sensitivity analyses assessed robustness and did not redefine the primary result after observing statistical significance. No analysis was interpreted as evidence of causal necessity or directed neural transmission.

### Ethics

The original study was approved by the Columbia University Institutional Review Board (Protocol AAAE3743), and all participants provided informed consent (Woo et al., 2015). The present reanalysis used publicly available, de-identified neuroimaging data from OpenNeuro (accession ds000140) and involved no new participant recruitment, data collection, or participant contact.

## Results

### Voluntary regulation shifted warmth/pain ratings in both directions

Stimulus intensity strongly predicted warmth/pain ratings (standardized beta = 0.698, 95% CI [0.658, 0.738], p < 0.001). After adjustment for stimulus intensity and task-position covariates, up-regulation increased ratings relative to passive trials (beta = 0.340, 95% CI [0.189, 0.491], p < 0.001), whereas down-regulation decreased them (beta = -0.289, 95% CI [-0.426, -0.151], p < 0.001; Figure 1A). Finite-cluster sensitivity preserved both standardized instruction effects (up: CR2 p < 0.001, wild-cluster p = 0.0001; down: CR2 p < 0.001, wild-cluster p < 0.0001; Supplementary Table S7a).

**Figure 1.**
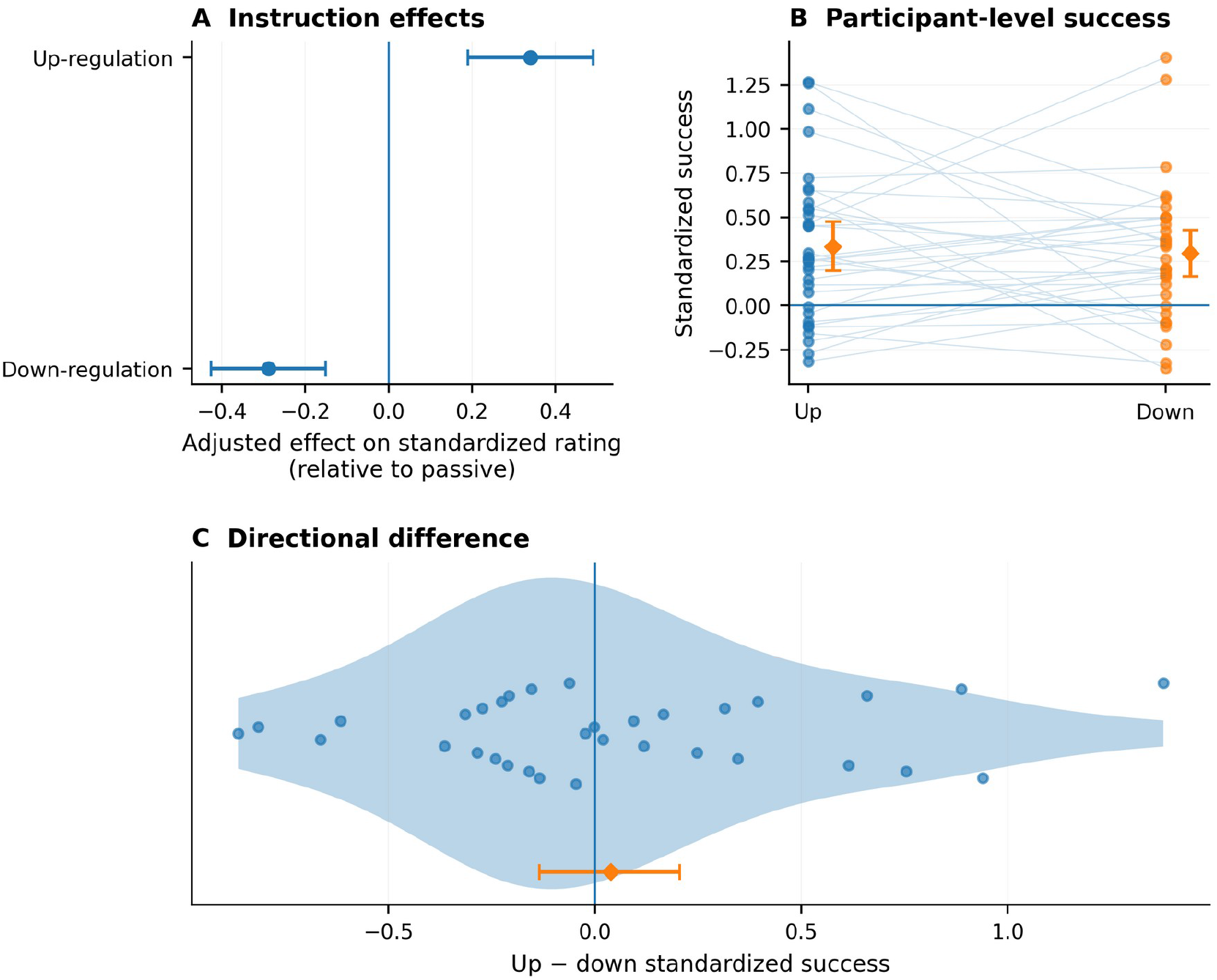
Behavioral validation of voluntary pain regulation. (A) Adjusted instruction effects on standardized ratings relative to passive trials; points show model estimates and whiskers show 95% confidence intervals, with zero indicating no instruction effect. (B) Participant-level direction-corrected standardized regulation success. Lines connect the same participant across up- and down-regulation; diamonds and whiskers show group means and bootstrap 95% confidence intervals. (C) Participant-level up-minus-down success differences. The violin summarizes the distribution, points show all 33 participants, and the diamond and whisker show the group mean and bootstrap 95% confidence interval. Both directions showed positive mean regulation success, whereas the mean directional difference was small and not reliable.

Mean direction-corrected standardized success was positive for up-regulation (M = 0.333, bootstrap 95% CI [0.196, 0.476], p < 0.001) and down-regulation (M = 0.295, bootstrap 95% CI [0.166, 0.427], p < 0.001; Figure 1B; Table 2). The mean up-minus-down difference was small and not reliable (M = 0.038, bootstrap 95% CI [-0.134, 0.206], p = 0.667; Figure 1C). The finite-cluster results were concordant: both direction-specific success means remained reliable (wild-cluster p ≤ 0.0002), whereas the up-minus-down asymmetry did not (CR2 p = 0.636; wild-cluster p = 0.632).

**Table 2.** Behavioral regulation effects and run-position checks.

| Analysis | Estimate | 95% CI | p |
| --- | --- | --- | --- |
| Up-regulation versus passive | $\beta = 0.340$ | [0.189, 0.491] | $< 0.001$ |
| Down-regulation versus passive | $\beta = -0.289$ | [-0.426, -0.151] | $< 0.001$ |
| Up-regulation success | $M = 0.333$ | [0.196, 0.476] | $< 0.001$ |
| Down-regulation success | $M = 0.295$ | [0.166, 0.427] | $< 0.001$ |
| Up-minus-down success | $M = 0.038$ | [-0.134, 0.206] | 0.667 |
| Down versus nearest passive | $M = -0.683$ | [-0.852, -0.513] | $< 0.001$ |
| Down versus following passive | $M = -0.248$ | [-0.411, -0.085] | 0.004 |
| Up versus nearest passive | $M = 0.117$ | [-0.064, 0.298] | 0.198 |
| Up versus following passive | $M = 0.369$ | [0.217, 0.520] | $< 0.001$ |

Note. Regression coefficients and participant mean effects are expressed on the within-participant standardized rating scale. Run-matched comparisons are participant-level standardized mean differences.

Because regulation runs occupied fixed positions, we compared regulation ratings with nearby passive runs. Down-regulation remained lower than the nearest passive comparator (M = -0.683, p < 0.001) and the following passive run (M = -0.248, p = 0.004). Up-regulation did not differ reliably from the nearest passive run (M = 0.117, p = 0.198) but was higher than the following passive run (M = 0.369, p < 0.001). Passive ratings declined across run position (slope = -1.19 rating units per run, p = 0.011), and the corresponding finite-cluster standardized slope was also negative (CR2 p = 0.029; wild-cluster p = 0.031), so condition-versus-passive effects retain a run-position constraint.

The standardized success score initially yielded a direction-by-expected-pain interaction (beta = -0.266, SE = 0.090, p = 0.003). The interaction was not reliable when success was expressed in raw units (beta = -0.191, p = 0.078), when observed raw ratings were analyzed directly (beta = -0.095, p = 0.289), or when success was normalized by the available room for change (beta = -0.0014, p = 0.671; Figure 2A). Boundary-trimmed analyses likewise weakened the pattern. In the finite-cluster model, the direction-by-expected-pain interaction was not reliable (CR2 p = 0.098; wild-cluster p = 0.097; Supplementary Table S7a).

**Figure 2.**
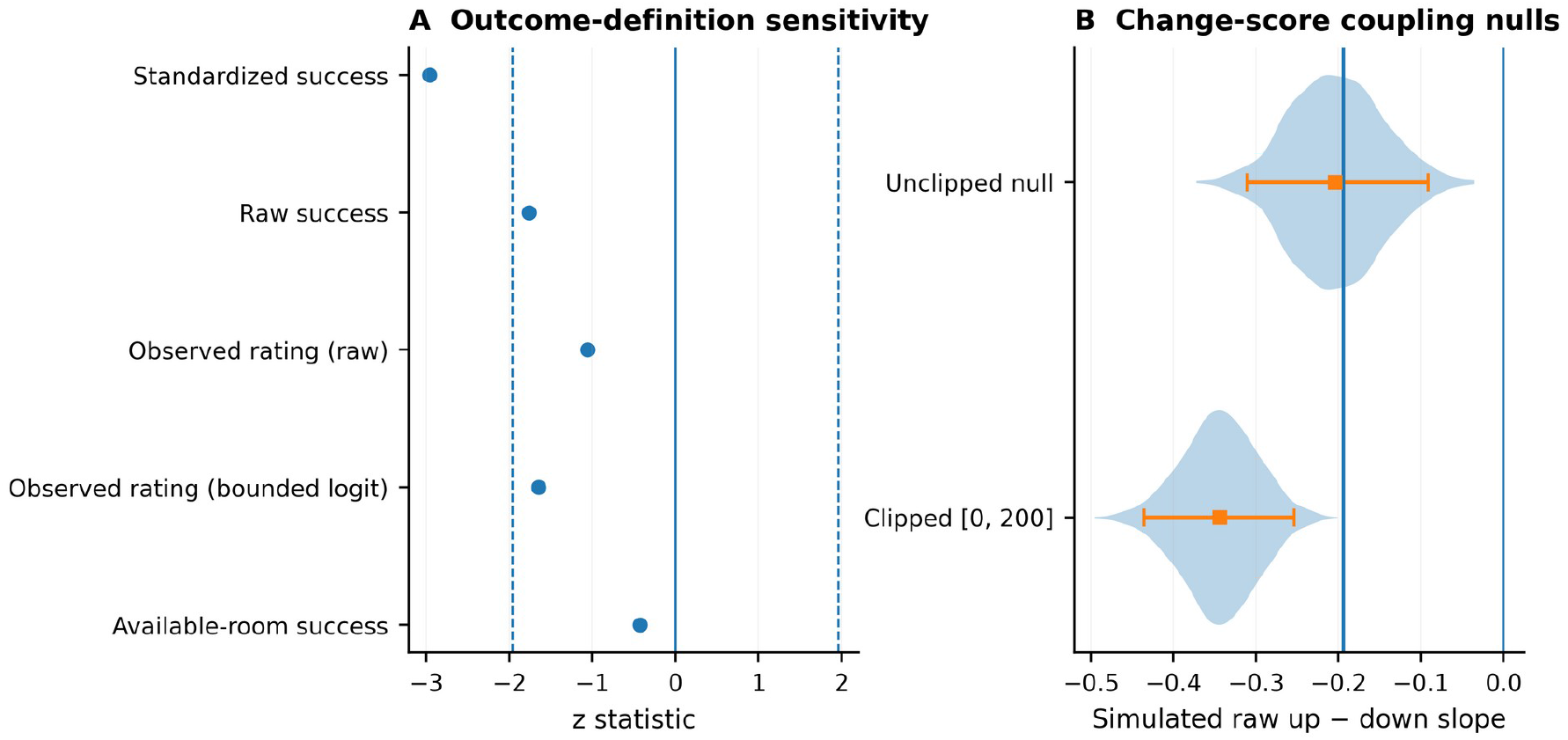
Sensitivity of the behavioral state-dependence result to outcome definition and change-score coupling. (A) z statistics for the direction-by-expected-pain interaction across alternative outcome definitions; dashed reference lines mark |z| = 1.96. Only the standardized-success definition survived the reported false-discovery-rate correction across the tested contrast family. (B) Simulated null distributions for the raw up-minus-down slope under unclipped and [0, 200]-clipped rating simulations. Violins show the null distributions, squares and whiskers show null medians and central 95% intervals, and the vertical line marks the observed raw slope. Under the unclipped null the observed value was close to the center of the distribution, whereas under the clipped null it lay beyond the simulated distribution.

Under the unclipped null, the observed raw slope (−0.194) was close to the null median (−0.204), lay at the 56.05th percentile and fell within the central 95% interval [−0.311, −0.092]. Clipping generated ratings to the observed 0–200 range shifted the simulated distribution further negative (median = −0.343; central 95% interval [−0.435, −0.254]); the observed slope was less negative than all 2,000 clipped-null values (Figure 2B). Thus, a negative direction-by-state slope could arise without a true interaction, and the apparent state dependence was not interpreted as evidence of a regulatory mechanism.

### Down-regulation was associated with regional response attenuation and left prefrontal engagement

Passive heat stimulation produced robust whole-brain responses, confirming sensitivity to nociceptive input. No voxels survived whole-brain FDR correction for up-minus-down, up-minus-passive, down-minus-passive or mean-regulation-minus-passive contrasts (Supplementary Figure S2; Supplementary Table S2).

Targeted participant-level fixed-effect maps identified seven mask-level down-minus-passive ROI effects after FDR correction across all 32 ROIs (Figure 3A; Table 3). Activity was lower in bilateral S2/operculum, left anterior and whole insula, and bilateral posterior insula. In the same contrast, left dlPFC/IFJ activity was higher (M = 0.0180, bootstrap 95% CI [0.0052, 0.0301], q = 0.0448). Because the left whole, anterior, and posterior insula masks overlap or are nested, the seven entries should not be interpreted as seven independent anatomical replications. The corrected regional pattern therefore describes a set of mask-level effects combining sensory and insular attenuation with increased activity in a lateral prefrontal control region.

**Figure 3.**
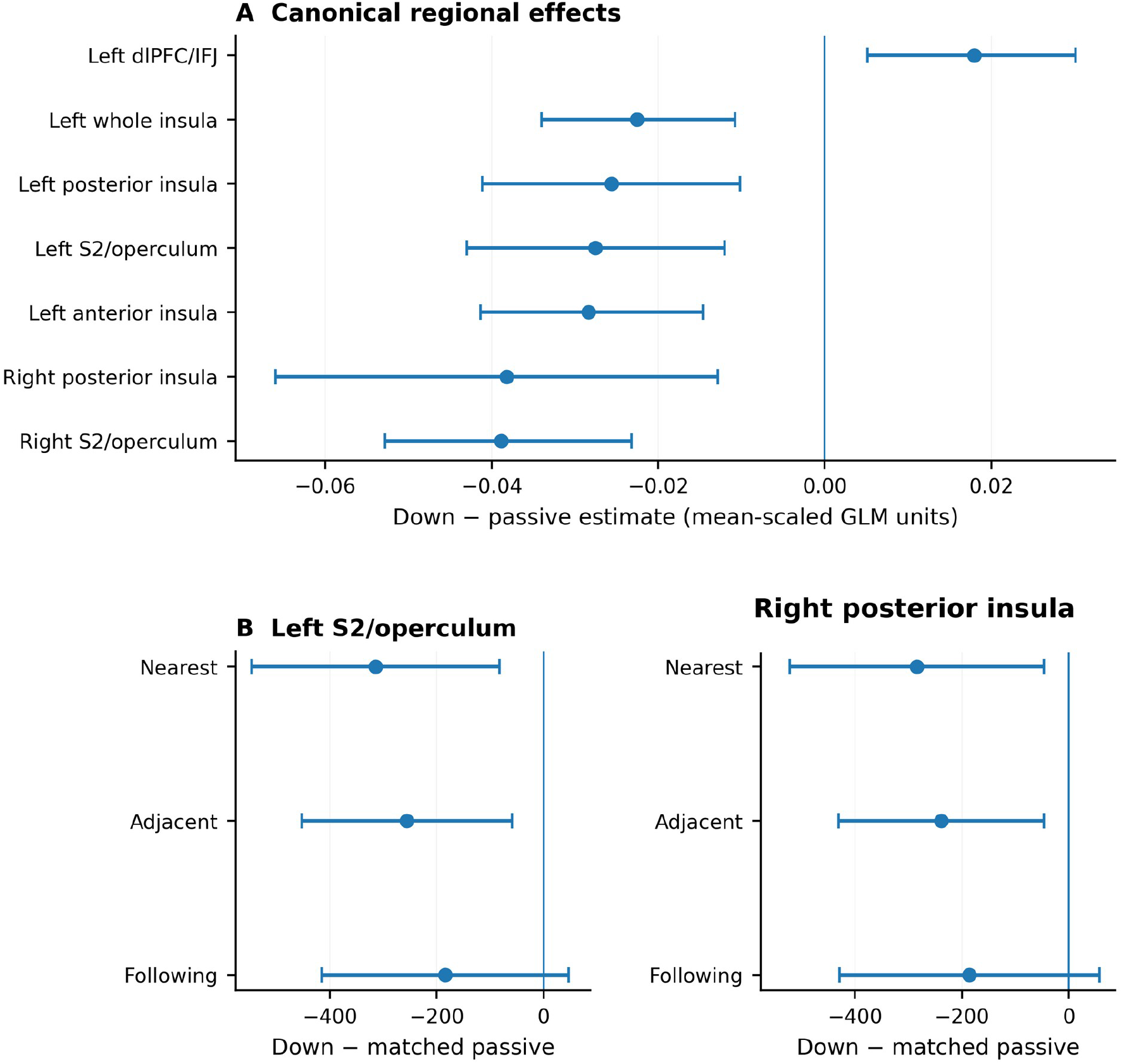
Regional down-regulation effects and run-position sensitivity. (A) Down-minus-passive effects for regions surviving false-discovery-rate correction across the 32-ROI family; points show mean effects and whiskers show 95% bootstrap confidence intervals. Negative values indicate lower regional responses during down-regulation, while the positive left dlPFC/IFJ estimate indicates greater activity. (B) Run-matched down-minus-passive sensitivity estimates for left S2/operculum and right posterior insula using nearest-passive, adjacent-mean-passive and following-passive comparators. Whiskers are 95% t intervals from participant-level paired differences. Nearest- and adjacent-passive estimates were nominally negative, but none of these selected-ROI comparisons survived the corresponding 32-ROI FDR correction. Panel A uses run-wise mean-scaled GLM contrast units, whereas Panel B uses native BOLD-intensity FWL-LSS units; magnitudes should not be compared directly across panels.

**Table 3.** Down-minus-passive ROI effects surviving FDR correction across all ROIs.

| ROI | Mean contrast | Bootstrap 95% CI | p | q |
| --- | --- | --- | --- | --- |
| Right S2/operculum | −0.0388 | [−0.0528, −0.0232] | < 0.001 | 0.00064 |
| Left anterior insula | −0.0284 | [−0.0414, −0.0146] | < 0.001 | 0.00508 |
| Left whole insula | −0.0225 | [−0.0340, −0.0108] | < 0.001 | 0.00840 |
| Left S2/operculum | −0.0276 | [−0.0430, −0.0120] | 0.0017 | 0.0134 |
| Left posterior insula | −0.0256 | [−0.0411, −0.0102] | 0.0030 | 0.0190 |
| Right posterior insula | −0.0382 | [−0.0659, −0.0128] | 0.0094 | 0.0448 |
| Left dlPFC/IFJ | 0.0180 | [0.0052, 0.0301] | 0.0098 | 0.0448 |

Note. Contrast estimates are participant-level fixed-effect ROI means from the run-wise mean-scaled GLM; confidence intervals are percentile bootstrap intervals from 5,000 participant-level resamples. The left whole, anterior, and posterior insula masks overlap or are nested and do not constitute independent anatomical replications.

The negative S2/opercular and left-insular estimates retained their direction under stricter run-level and participant-level motion exclusions; the strongest bilateral S2/opercular and left-insular effects also remained FDR-significant in at least one sensitivity set (Supplementary Figure S4; Supplementary Table S4a). The positive dlPFC/IFJ estimate retained its sign but did not survive all-ROI correction in the motion-exclusion sets. Run-position-matched analyses comprised eight regulation-direction-by-comparator families, with Benjamini–Hochberg correction across all 32 ROI features within each family. Figure 3B and Supplementary Table S4b display two selected down-regulation ROIs for three comparator definitions: uncorrected p values ranged from 0.009 to 0.130, whereas the corresponding 32-ROI-family q values ranged from 0.224 to 0.829; none survived correction, and effects were not reliable against the following passive run. Full results for all 256 tests are provided in Supplementary Data S1. The displayed run-matched estimates use native-intensity LSS units and are not numerically comparable with the mean-scaled GLM contrasts in Figure 3A. The design therefore does not completely separate condition, run position, and task-context components.

NPS expression depended on the estimation path. In participant-level contrast maps, down-minus-passive NPS was negative (M = −0.00484, bootstrap 95% CI [−0.00697, −0.00278], q = 0.00024). In the rating-adjusted FWL-LSS path, the corresponding weighted-mean contrast was not reliable (M = −3.34, 95% CI [−23.26, 16.57], t(32) = −0.34, p = 0.735, q = 0.735). Up-minus-passive, mean-regulation-minus-passive, and up-minus-down FWL-LSS contrasts were also not reliable (all p ≥ 0.215). By contrast, participant-specific passive temperature slopes were strongly positive (M = 54.32, 95% CI [43.40, 65.23], t(32) = 10.14, p < 0.001, q < 0.001), and trial-level participant-fixed-effects models likewise showed a strong temperature coefficient without condition effects. NPS therefore validated thermal encoding but did not provide method-independent evidence for regulation-related attenuation.

### Up-regulation success covaried with an exploratory lagged prefrontal–valuation composite

The pathway had been selected in the archived 116-correlation screen (screening q = 0.0127). After rating-period adjustment and FWL re-estimation, standardized up-regulation success remained associated with the selected lagged left dlPFC/IFJ–valuation-region composite (r = −0.546, p = 0.0010, bootstrap 95% CI [−0.764, −0.270]; Figure 4A; Table 4). The association was similar using Spearman correlation (ρ = −0.549, p < 0.001) and remained negative after adjustment for regulation order, passive mean rating, and passive sensory slope (standardized β = −0.464, HC3 p = 0.0023; bootstrap 95% CI [−0.681, −0.194]; label-permutation p = 0.0007). All 33 leave-one-participant-out estimates retained the negative sign and ranged from r = −0.604 to r = −0.467. The updated score was highly correlated with the archived selected score (r = 0.985), indicating that rating adjustment changed the magnitude modestly rather than changing the participant ordering.

**Figure 4.**
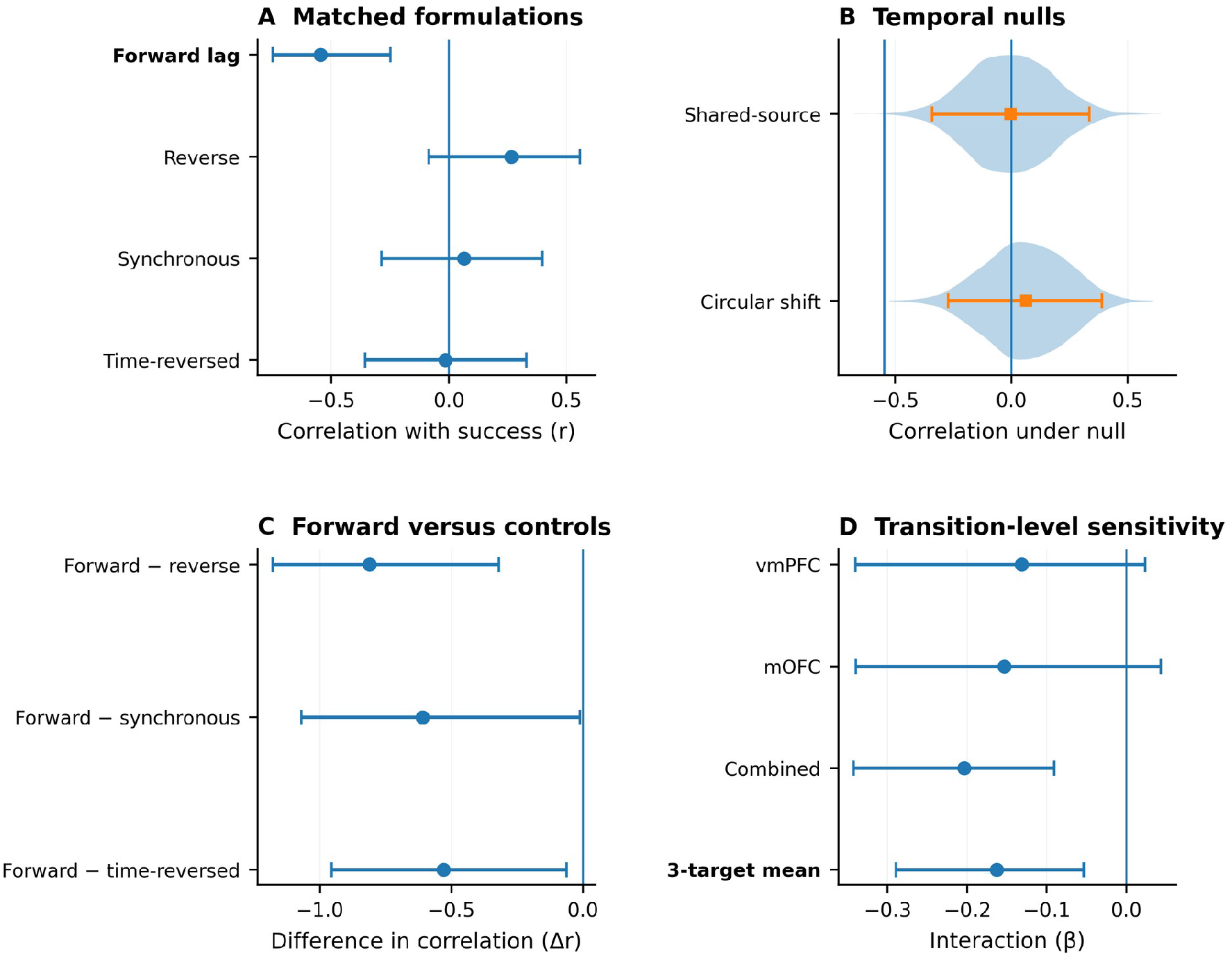
Rating-adjusted exploratory lagged prefrontal-valuation coupling, matched controls and transition-level sensitivity. (A) Correlations between up-regulation success and the selected forward composite, reverse-direction, synchronous zero-lag and time-reversed formulations, with descriptive Fisher-z 95% intervals. (B) Shared-source permutation and nonzero within-run circular-shift null distributions; the vertical line marks the observed forward correlation. (C) Participant-bootstrap differences between the forward association and matched controls; the forward formulation was more negative than each matched control in the reported bootstrap comparisons, with FDR correction applied across the three contrasts. (D) Covariate-adjusted transition-level sensitivity using all 297 up-regulation transitions from the archived rating-adjusted AR(1)-prewhitened trial estimates. Points show success-by-lagged-source interaction coefficients for vmPFC, mOFC, the combined vmPFC/mOFC target and their equal-weight three-target mean; whiskers are participant-cluster bootstrap 95% intervals. All coupling follow-ups remain conditional on the same-sample selected pathway and do not constitute independent validation.

**Table 4.** Exploratory lagged prefrontal–valuation coupling and matched controls.

| Analysis | Estimate | 95% CI / null interval | p | Control / correction |
| --- | --- | --- | --- | --- |
| Selected exploratory forward composite | $r = -0.546$ | $[-0.764, -0.270]$ | 0.0010 | archived screening $q = 0.0127$ |
| Covariate-adjusted model | $\beta = -0.464$ | $[-0.681, -0.194]$ | 0.0023 | label-permutation $p = 0.0007$ |
| Reverse-direction composite | $r = 0.265$ | $[-0.086, 0.558]$ | 0.136 | matched control; Fisher-z CI |
| Synchronous zero-lag composite | $r = 0.064$ | $[-0.286, 0.398]$ | 0.725 | matched control; Fisher-z CI |
| Time-reversed composite | $r = -0.016$ | $[-0.357, 0.329]$ | 0.930 | matched control; Fisher-z CI |
| Shared-source permutation null | $r = -0.546$ | null $[-0.341, 0.336]$ | 0.0014 | 5,000 iterations |
| Nonzero circular-shift null | $r = -0.546$ | null $[-0.271, 0.389]$ | 0.0014 | 5,000 iterations |
| Forward minus reverse | $\Delta r = -0.811$ | $[-1.178, -0.320]$ | 0.0022 | $q = 0.0066$ |
| Forward minus zero-lag | $\Delta r = -0.609$ | $[-1.070, -0.011]$ | 0.0470 | $q = 0.0470$ |
| Forward minus time-reversed | $\Delta r = -0.530$ | $[-0.954, -0.063]$ | 0.0258 | $q = 0.0387$ |

Note. The pathway and outcome association was selected in the same sample. Rating-adjusted matched-control, permutation, circular-shift, and bootstrap analyses are conditional post-selection characterizations and do not repeat or correct the 116-correlation screening stage.

Matched reverse-direction (r = 0.265, p = 0.136), synchronous zero-lag (r = 0.064, p = 0.725), and time-reversed (r = −0.016, p = 0.930) composites were not related to success. The observed forward correlation fell outside both the shared-source within-run permutation null (empirical two-sided p = 0.0014; central 95% interval [−0.341, 0.336]; Figure 4B) and the nonzero circular-shift null (p = 0.0014; interval [−0.271, 0.389]). Direct bootstrap differences favored the forward formulation over reverse direction (Δr = −0.811, 95% CI [−1.178, −0.320], q = 0.0066), synchronous coupling (Δr = −0.609, 95% CI [−1.070, −0.011], q = 0.0470), and time reversal (Δr = −0.530, 95% CI [−0.954, −0.063], q = 0.0387; Figure 4C; Table 4). These analyses characterize the selected statistical association but do not correct or independently validate the preceding screening stage.

AR(1) prewhitening yielded a closely corresponding forward score (r = 0.963 with the primary forward score) and preserved its association with up-regulation success (r = -0.585, p = 0.00035; Supplementary Table S7b). Reverse-direction (r = 0.242, p = 0.175), synchronous zero-lag (r = 0.097, p = 0.591), and time-reversed (r = 0.010, p = 0.956) controls remained unrelated to success. The forward effect fell outside the shared-source permutation (empirical two-sided p = 0.0008) and nonzero circular-shift (p = 0.0014) null distributions, and all three direct forward-minus-control differences remained FDR-significant (all q ≤ 0.021). These temporal-noise checks remain conditional on the selected pathway and do not repeat the screening stage.

A transition-level sensitivity analysis reduced the dependence of inference on the nine-transition participant-specific coefficients. Across all 297 up-regulation transitions, the equal-weight mean success-by-source interaction across vmPFC, mOFC, and combined vmPFC/mOFC targets was negative in the unadjusted model (beta = -0.121; participant-label permutation p = 0.0136) and became more negative after adjustment for regulation order, passive mean rating, and passive sensory slope (beta = -0.163; participant-label permutation p = 0.0003). The adjusted effect remained outside a Freedman-Lane conditional permutation null (p = 0.0102), and its participant-cluster bootstrap 95% CI excluded zero [-0.290, -0.054]. All 33 leave-one-participant-out adjusted estimates remained negative (range, -0.223 to -0.134). Target-specific effects were strongest for vmPFC and the combined vmPFC/mOFC definition, whereas mOFC alone was less stable. Random-slope mixed models yielded coefficients in the same direction but showed convergence or boundary warnings for two of the three target definitions. These analyses used the archived rating-adjusted AR(1)-prewhitened trial estimates and remain conditional on the selected pathway.

Split-half reliability was not estimated because each half would contain only four or five transitions for a four-parameter model. Each participant-direction coefficient was therefore still estimated from only nine transitions and five residual degrees of freedom, leaving individual-level reliability unknown despite the group-level transition sensitivity. Across six corrected session-wise DCM variants, no consistent group-average B-matrix effect was retained for up-regulation versus passive stimulation or down-regulation versus passive stimulation, and no stable family-wide up-minus-down pattern emerged. The DCM family also did not identify a robust up-specific success effect corresponding to the selected lagged dlPFC/IFJ–valuation association. One model-specific up-minus-down effect was retained for PAG-to-hypothalamus modulation in the right-amygdala affective variant, but it was absent from the homologous left-amygdala model and was treated as exploratory. Across models, individual differences were more consistently associated with passive rating level, passive sensory slope, and regulation order. Thus, DCM did not independently validate an up-specific directed mechanism for the trial-wise coupling result.

## Discussion

Voluntary regulation reliably changed warmth/pain ratings in both instructed directions, and the strongest neural associations emerged at complementary analytical levels. During down-regulation, left dlPFC/IFJ activity increased while bilateral S2/opercular and insular responses decreased relative to passive stimulation. Across participants, up-regulation success was associated with an exploratory, post-selection lagged dlPFC/IFJ-vmPFC/mOFC composite that remained evident after rating-period adjustment. These findings do not establish separate mechanisms for increasing and decreasing pain; they identify a condition-level regional pattern and an individual-difference coupling association, while the corrected DCM family provided no consistent mechanistic convergence.

The down-regulation pattern combined greater activity in a lateral prefrontal region implicated in cognitive control and pain reappraisal (Wiech et al., 2006) with lower responses in cortical regions implicated in sensory-discriminative and nociceptive-affective processing (Petre et al., 2022; Tracey and Mantyh, 2007). All seven mask-level entries survived correction across 32 ROIs, and negative S2/opercular and insular estimates retained their direction under motion-sensitivity analyses. These entries are not seven independent anatomical replications because the left whole, anterior, and posterior insula masks overlap or are nested. The joint pattern is compatible with prefrontal engagement accompanying attenuation of regional sensory and insular responses rather than a reduction in a pain-specific representation. Meta-analytic evidence likewise supports multiple neural routes for cognitive and emotional pain modulation (Modernell et al., 2026). Spatial generality is limited by the absence of corrected whole-brain regulation effects and by correction being performed within, rather than across, ten ROI effect families. Run-matched comparisons provided only uncorrected support for selected sensory regions. Contrast-map NPS attenuation also did not replicate in the fully rating-adjusted FWL-LSS trial-wise path, despite strong temperature sensitivity. The most defensible interpretation is therefore a down-regulation-associated regional pattern that may contain run-position and task-context components, without method-independent multivariate-signature support.

The up-regulation result captured variation between participants rather than an average condition contrast. The rating-adjusted lagged prefrontal-valuation composite remained associated with success after covariate adjustment and leave-one-participant-out analyses. Expectation studies similarly suggest that amplification and attenuation can differ in temporal and regional expression, although expectation paradigms are not equivalent to voluntary regulation (Mourkojannis et al., 2025; Wittkamp et al., 2024; Wolf et al., 2024). The tested forward-lagged association was more negative than matched reverse-direction, synchronous zero-lag, and time-reversed formulations and fell outside shared-source permutation and circular-shift null distributions. AR(1) prewhitening preserved the pattern, and direct modeling of all 297 up-regulation transitions retained a negative success-by-lagged-source interaction after adjustment for regulation order, passive mean rating, and passive sensory slope. These analyses reduce concern that the group association arose solely from highly uncertain participant-specific coefficients, but they do not provide independent validation or establish physiological directionality. The pathway was selected in the same sample, its valuation masks overlapped, and each participant contributed only nine within-run transitions to a four-parameter model. The negative coefficient should not be interpreted as inhibitory influence, signal transmission, or effective connectivity. Across six run-separated DCM variants, neither regulation-versus-passive contrasts nor up-specific success effects consistently converged with the lagged association. The result is therefore a post-selection candidate prefrontal-valuation coupling pattern for confirmatory testing. Related systems-level work supports distributed prefrontal involvement in regulation (Bo et al., 2024; He et al., 2023; Wang et al., 2026; Zhang et al., 2025), while systematic-review evidence cautions that individual-difference associations can vary across studies and analytical choices (Morawetz and Basten, 2024).

The behavioral analyses add a methodological point. Standardized success initially suggested direction-dependent regulation as a function of passive-expected pain, but the interaction weakened or disappeared with alternative outcome definitions or direct analysis of observed ratings. Similar negative slopes arose under change-score coupling nulls without a true direction-by-state effect, and clipping simulated ratings to the bounded scale shifted the null further negative. This follows from using expected pain in both the predictor and a bounded change-score outcome. It does not diminish the clear instruction effects; rather, state-dependent claims require outcome definitions that separate psychological modulation from arithmetic coupling and rating-scale limits (Bland and Altman, 1994; Oldham, 1962; Tu and Gilthorpe, 2007).

The findings extend the original ds000140 account rather than overturning it. Woo et al. (2015) reported strong stimulus-intensity effects on NPS, no reliable canonical up-versus-down NPS effect, and a vmPFC-nucleus accumbens mediation pathway for self-regulation. We likewise observed robust heat-related NPS expression and no reliable up-versus-down NPS effect in the corrected trial-wise estimator. The direction-resolved analyses add prefrontal engagement with S2/opercular-insular attenuation during down-regulation and an individual-difference association between up-regulation success and lagged lateral-prefrontal-valuation coupling. The absence of consistent DCM regulation-versus-passive effects indicates that the coupling result should not be treated as a model-based estimate of the original mediation pathway. Because these analyses use different contrasts and inferential levels, they are complementary rather than direct replications or contradictions. Although this study is non-clinical, evidence that emotion-regulation-focused interventions can improve pain-related outcomes supports the broader relevance of voluntary regulation without implying that the present neural associations mediate treatment benefit (Norman-Nott et al., 2024).

Several limitations define the scope of the conclusions. The sample included 33 participants, with one up-regulation and one down-regulation run per participant; finite-cluster sensitivities supported the primary behavioral conclusions, but inference still rests on 33 independent clusters. Condition, run position, and order were structurally linked, preventing complete separation of instruction effects from session context. Targeted analyses were not preregistered; ROI FDR correction was performed within ten effect families, and reported left-insula entries include overlapping or nested masks. The coupling composite was selected in the same sample, combined related target definitions, and lacked independent validation. Participant-level coupling coefficients were based on nine transitions and five residual degrees of freedom, leaving individual-level reliability unknown. Transition-level clustered sensitivity reduced dependence on those slopes but did not address same-sample selection. fMRI temporal resolution and hemodynamic variability limit directional interpretation of lagged regressions; AR(1), permutation, and circular-shift controls do not capture every feature of trial-wise fMRI estimates or repeat the screening stage. The released outcome combines warmth and pain rather than isolating pain. The released temperature field (40.8–47.3) differs from the nominal protocol temperatures reported originally (44.3–49.3°C), and no conversion is documented. The demographic field encoded 22 participants as female and 11 as male, but source metadata did not document how sex was ascertained and gender identity was unavailable; the sample was not designed or powered for sex-stratified inference. NPS results differed across estimation paths, and the trial-wise estimator should not be treated as an exact reconstruction of the original pipeline. Finally, the six DCM variants were exploratory and yielded no consistent regulation-versus-passive pattern. Independent, preregistered studies with denser within-participant sampling are needed.

In conclusion, voluntary regulation shifted warmth/pain ratings in both directions and showed complementary direction-sensitive neural correlates. Down-regulation combined greater left dlPFC/IFJ activity with lower S2/opercular and insular responses, whereas up-regulation success covaried with a post-selection candidate lagged coupling pattern between lateral prefrontal control and vmPFC/mOFC valuation regions. Temporal controls, AR(1) prewhitening, and transition-level sensitivity supported the statistical association, but same-sample selection, uncertain individual-level reliability, and lack of consistent DCM convergence limit mechanistic interpretation.

## Acknowledgments

We thank Tor D. Wager for providing access to the Neurologic Pain Signature weight map used in this study. We also thank the investigators of the original study for making ds000140 publicly available through OpenNeuro.

## Declarations

### Data and code availability

The original neuroimaging and behavioral data are publicly available from OpenNeuro (ds000140, version 00001) (Woo et al., 2018). The analysis code, derived tabular results, machine-readable source tables, quality-control summaries, final reporting assets, and transition-level sensitivity analysis are publicly available through GitHub repository version 1.1.0 (Kharghani and Gholami, 2026) and an immutable versioned Zenodo archive at https://doi.org/10.5281/zenodo.23010861. Large imaging derivatives, ROI masks, and MATLAB DCM objects are not redistributed but can be regenerated from the public dataset using the documented workflows. The NPS weight map is not included and must be obtained independently from its original source.

### Generative AI disclosure

OpenAI ChatGPT was used to assist with language editing, code drafting and debugging for additional sensitivity analyses, preparation and refinement of data visualizations, and manuscript/document formatting. All AI-assisted code, numerical results, statistical outputs, references, figures, and manuscript text were independently reviewed and verified by the authors against the source data and analysis outputs. The authors made all scientific and interpretive decisions and take full responsibility for the final work.

### Author contributions

Meysam Kharghani: Conceptualization, Methodology, Software, Formal analysis, Investigation, Data curation, Validation, Visualization, Project administration, Supervision, Writing - original draft, Writing - review and editing. Elahe Gholami: Resources, Data curation, Formal analysis, Visualization, Writing - review and editing. Both authors read and approved the final manuscript.

