## Supplementary Information for "Direction-sensitive neural correlates of voluntary pain regulation: regional response attenuation during down-regulation and exploratory prefrontal-valuation coupling"

This Supplementary Information provides quality-control, whole-brain, state-dependence, run-position, regional, Neurologic Pain Signature (NPS), exploratory lagged-coupling, finite-cluster inference, AR(1) temporal-noise sensitivity, transition-level coupling sensitivity, and dynamic-causal-modeling analyses that complement the main manuscript. The analysis code, complete machine-readable results, transition-level sensitivity outputs, final reporting source tables, and reproducibility materials correspond to repository version 1.1.0 and are archived in the associated versioned Zenodo record (https://doi.org/10.5281/zenodo.23010861).

### Supplementary Note 1. Quality control and whole-brain inference

The released dataset contained 297 functional runs from 33 participants. The main imaging analyses retained 288 runs (222 passive, 33 up-regulation and 33 down-regulation) and excluded nine passive runs. A run was excluded if at least one documented severe-motion criterion was met: mean framewise displacement (FD) greater than 0.50 mm, more than 30% of volumes with FD greater than 0.5 mm, more than 10% of volumes with FD greater than 1.0 mm, or maximum FD greater than 5.0 mm. The availability and length-consistency criteria did not exclude any observed run. All 66 regulation runs were retained. Visual inspection indicated acceptable brain coverage, spatial alignment and preprocessing quality in the retained data.

Passive heat stimulation survived voxelwise false-discovery-rate correction, whereas no regulation-related whole-brain contrast—up minus down, up minus passive, down minus passive or mean regulation minus passive—contained corrected voxels. These results establish sensitivity to nociceptive stimulation but do not provide corrected whole-brain corroboration of the targeted regional, signature or pathway findings.

Passive ratings declined across run position (raw slope = −1.19 rating units per run, $p = 0.0109$). Down-regulation remained lower than the nearest and following passive comparators, whereas up-regulation was not reliably higher than the nearest passive run but was higher than the following passive run. Regulation-versus-passive contrasts are therefore interpreted as condition-level effects that may include run-position and task-context components.

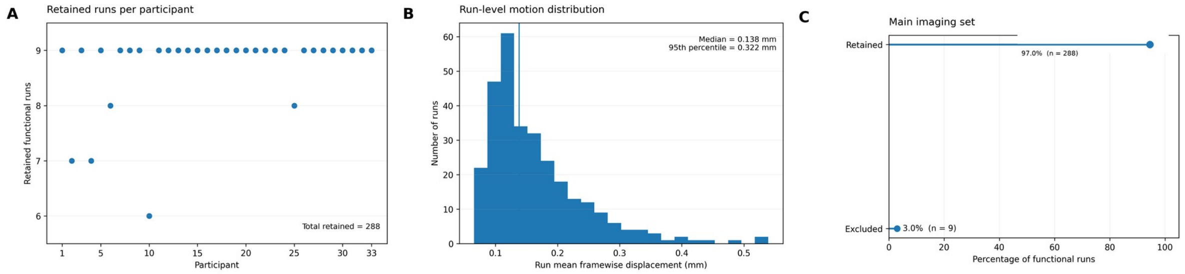


Supplementary Figure 1. Imaging quality control. (A) Number of retained functional runs per participant. (B) Distribution of run mean framewise displacement; the vertical line marks the median. (C) Percentage and count of functional runs retained and excluded from the main imaging set.

**Supplementary Table S1. Primary imaging data retained after quality control**

| Condition | Runs acquired | Runs retained | Runs excluded | Participants retained | Mean FD, mm |
|---|---|---|---|---|---|
| Passive | 231 | 222 | 9 | 33 | 0.156 |
| Up-regulation | 33 | 33 | 0 | 33 | 0.176 |
| Down-regulation | 33 | 33 | 0 | 33 | 0.157 |
| Total | 297 | 288 | 9 | 33 | 0.158 |

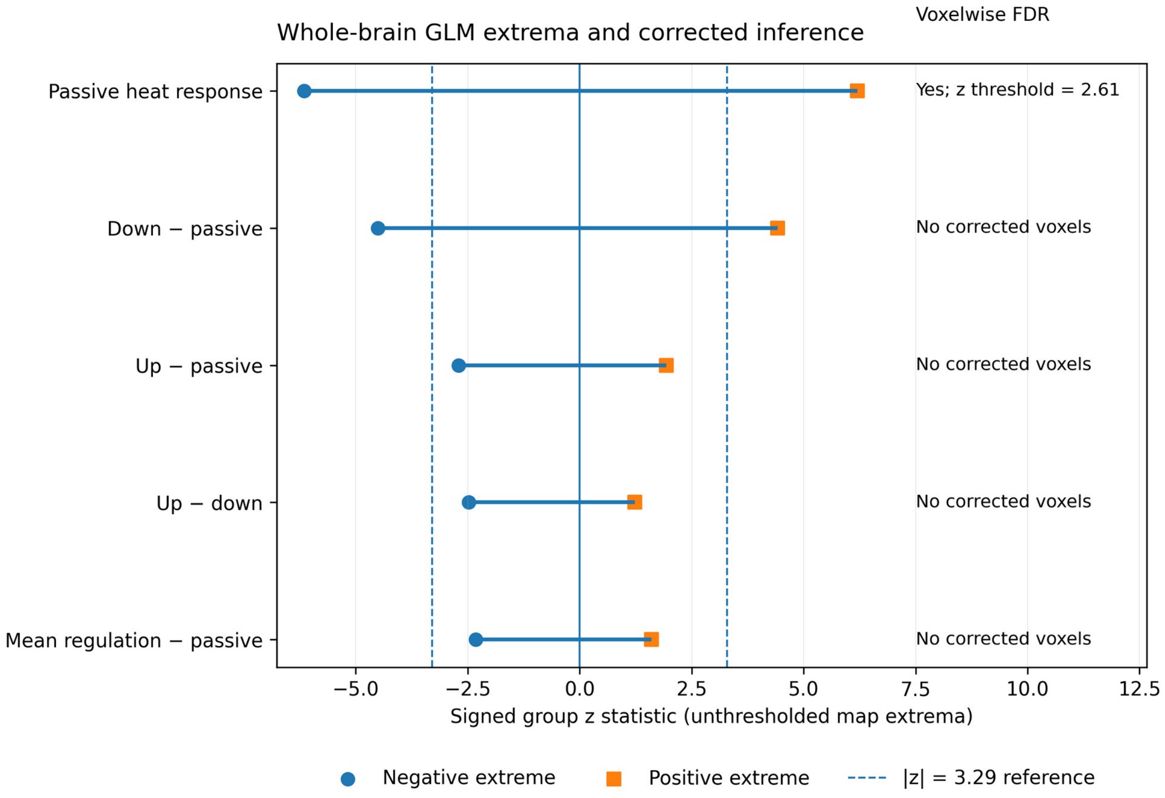


Supplementary Figure 2. Whole-brain general linear model summary. Horizontal segments show the signed minimum-to-maximum group z-statistic range in each unthresholded main-analysis map; circles and squares mark the negative and positive extrema, respectively. Dashed vertical lines at |z| = 3.29 are descriptive reference lines only. Passive heat stimulation contained voxels surviving voxelwise false-discovery-rate correction (FDR z threshold = 2.61), whereas none of the regulation-related whole-brain contrasts contained FDR-corrected voxels. Extrema summarize map magnitude and do not localize effects spatially.

**Supplementary Table S2. Whole-brain general linear model summary**

| Contrast | n | FDR-surviving voxels | FDR z threshold | Minimum z | Maximum z | Maximum \|z\| |
|---|---|---|---|---|---|---|
| Passive heat response | 33 | Yes | 2.611 | -6.148 | 6.205 | 6.205 |
| Down – passive | 33 | No | — | -4.508 | 4.418 | 4.508 |
| Up – passive | 33 | No | — | -2.702 | 1.935 | 2.702 |
| Up – down | 33 | No | — | -2.476 | 1.232 | 2.476 |
| Mean regulation – passive | 33 | No | — | -2.316 | 1.604 | 2.316 |

### Supplementary Note 2. State-dependence sensitivity

The main standardized success score produced a negative direction-by-expected-pain interaction. The interaction weakened when the outcome was expressed as raw success, observed ratings, bounded-logit ratings or available-room-normalized success. Cross-fitting preserved the interaction for standardized success and for some bounded formulations, but not for raw success or observed raw ratings. The non-cross-fitted fractional-logit fit was numerically non-interpretable: coefficients were on the order of 10^14–10^15 and associated standard errors or p values were non-finite, a pattern consistent with separation or numerical failure.

Coupling-null simulations tested whether the raw interaction could arise without a true direction-by-state effect. In 2,000 unbounded simulations, the null mean was −0.202, the median was −0.204 and the central 95% interval was [−0.311, −0.092]. The observed slope of −0.194 lay at the 56.05th percentile. Clipping generated ratings to the observed 0–200 range shifted the distribution further negative (mean = −0.344; median = −0.343; central 95% interval [−0.435, −0.254]); the observed slope was less negative than all clipped-null values. Because neither null distribution was centered at zero, absolute-value two-sided p values were not interpreted. The apparent baseline-state pattern is therefore considered sensitive to change-score geometry and scale boundaries rather than evidence of a regulatory mechanism.

**Supplementary Figure 3**

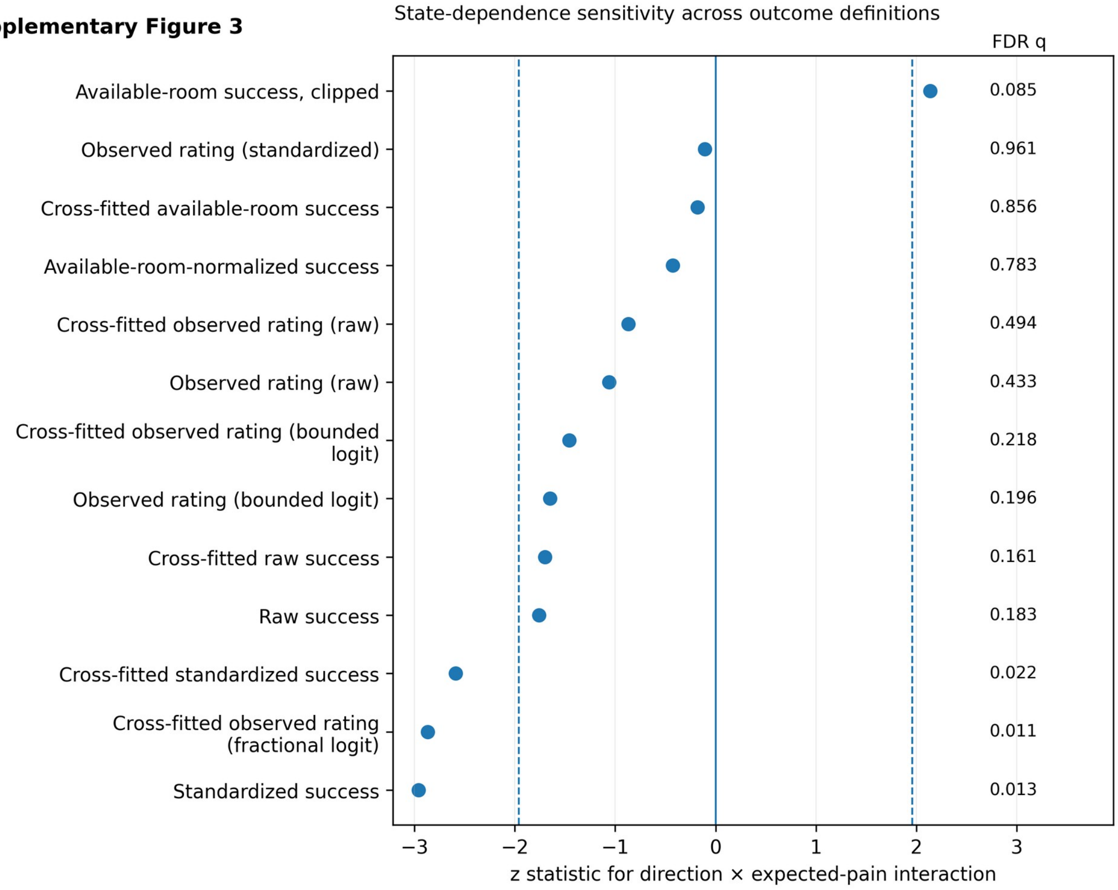


Supplementary Figure 3. State-dependence sensitivity across interpretable outcome definitions. Points show z statistics for the direction-by-expected-pain interaction; dashed lines mark |z| = 1.96 and q values show the reported false-discovery-rate correction. The numerically unstable non-cross-fitted fractional-logit model is omitted from the figure and remains documented in Supplementary Table S3.

**Supplementary Table S3. Direction-by-expected-pain interaction across outcome definitions**

| Outcome/ model | Estimate | SE | z | p | FDR q | Diagnostic note |
|---|---|---|---|---|---|---|
| Standardized success | -0.2655 | 0.0898 | -2.958 | 0.003 | 0.013 | Non-cross-fitted; q across all reported contrasts |

| Outcome/ model | Estimate | SE | z | p | FDR q | Diagnostic note |
|---|---|---|---|---|---|---|
| Raw success | -0.1915 | 0.1087 | -1.761 | 0.078 | 0.183 | Non-cross-fitted ; q across all reported contrasts |
| Observed rating (standardized) | -0.0079 | 0.0744 | -0.107 | 0.915 | 0.961 | Non-cross-fitted ; q across all reported contrasts |
| Observed rating (raw) | -0.0948 | 0.0893 | -1.061 | 0.289 | 0.433 | Non-cross-fitted ; q across all reported contrasts |
| Observed rating (bounded logit) | -1.3180 | 0.7997 | -1.648 | 0.099 | 0.196 | Non-cross-fitted ; q across all reported contrasts |
| Observed rating (fractional logit; unstable) | Not interpretable | — | — | — | — | Coefficients on the order of 10^14–10^15 with non-finite standard errors or p values; excluded from inference |
| Available-room-normalized success | -0.0014 | 0.0033 | -0.425 | 0.671 | 0.783 | Non-cross-fitted ; q across all reported contrasts |
| Available-room-normalized success (clipped) | 0.0027 | 0.0012 | 2.138 | 0.033 | 0.085 | Non-cross-fitted ; q across all reported contrasts |
| Cross-fitted standardized success | -0.2634 | 0.1017 | -2.590 | 0.010 | 0.022 | Cross-fitted passive baseline; q across model terms |
| Cross-fitted raw success | -0.1943 | 0.1143 | -1.699 | 0.089 | 0.161 | Cross-fitted passive baseline; q across model terms |
| Cross-fitted observed rating (raw) | -0.0812 | 0.0933 | -0.870 | 0.384 | 0.494 | Cross-fitted passive baseline; q across model terms |
| Cross-fitted observed rating (bounded logit) | -1.2189 | 0.8368 | -1.457 | 0.145 | 0.218 | Cross-fitted passive baseline; q across model terms |
| Cross-fitted observed rating (fractional logit) | -1.4692 | 0.5124 | -2.868 | 0.004 | 0.011 | Cross-fitted passive baseline; q across model terms |
| Cross-fitted available-room success | -0.0007 | 0.0037 | -0.182 | 0.856 | 0.856 | Cross-fitted passive baseline; q across model terms |

### Supplementary Note 3. Regional, run-matched and NPS analyses

Within the focal down-minus-passive contrast family, seven mask-level regional effects survived Benjamini–Hochberg false-discovery-rate correction across 32 ROI features. Six estimates were negative—bilateral S2/operculum, left anterior and whole insula, and bilateral posterior insula—and left dlPFC/IFJ was positive. These seven mask-level results are not seven anatomically independent replications: the left whole-insula, anterior-insula, and posterior-insula masks overlap or are nested. Across the complete ROI analysis, 10 contrast/effect families were evaluated; FDR was controlled across 32 ROI features separately within each family, not across all 10 families. The down-minus-passive family was emphasized in the main text; the remaining families should be interpreted as secondary or exploratory.

Run-matched ROI analyses compared each regulation run with the nearest passive run, the mean of adjacent passive runs, and the following passive run. Multiplicity correction was not limited to the six displayed cells. It comprised eight direction-by-comparator families (two regulation directions × four passive comparators), with Benjamini–Hochberg correction across all 32 ROI features within each family. For the six selected down-regulation estimates displayed in Supplementary Table S4b, uncorrected p values ranged from 0.009 to 0.130 and corresponding within-family q values ranged from 0.224 to 0.829; none survived $q < 0.05$. The full set of 256 tests is provided in Supplementary Data S1. These FWL-LSS estimates are differences in unscaled native fMRIPrep BOLD-intensity units and are not numerically comparable with the run-wise mean-scaled GLM contrast estimates in Figure 3A and Supplementary Table S4a.

NPS findings depended on estimation path. Direct participant contrast maps showed lower down-minus-passive NPS expression ($M = -0.00484$, 95% CI [−0.00697, −0.00278], $p < 0.001$), which was similar after excluding one influential participant ($M = -0.00460$). The corrected trial-wise analysis used rating-adjusted FWL-LSS on 6-mm-smoothed BOLD data from all 288 retained runs and yielded 3,102 stimulation-trial estimates. Its weighted-mean down-minus-passive contrast was not reliable ($M = -3.34$, 95% CI [−23.26, 16.57], $t(32) = -0.34$, $p = 0.735$, $q = 0.735$). Up-minus-passive, mean-regulation-minus-passive, and up-minus-down contrasts were also not reliable. Passive temperature slopes were strongly positive ($M = 54.32$, 95% CI [43.40, 65.23], $t(32) = 10.14$, $p < 0.001$, $q < 0.001$), and trial-level participant-fixed-effects models likewise retained a strong temperature coefficient without reliable condition effects. Thus, the corrected trial-wise estimator validated thermal encoding but did not support regulation-related NPS attenuation.

**Supplementary Figure 4**

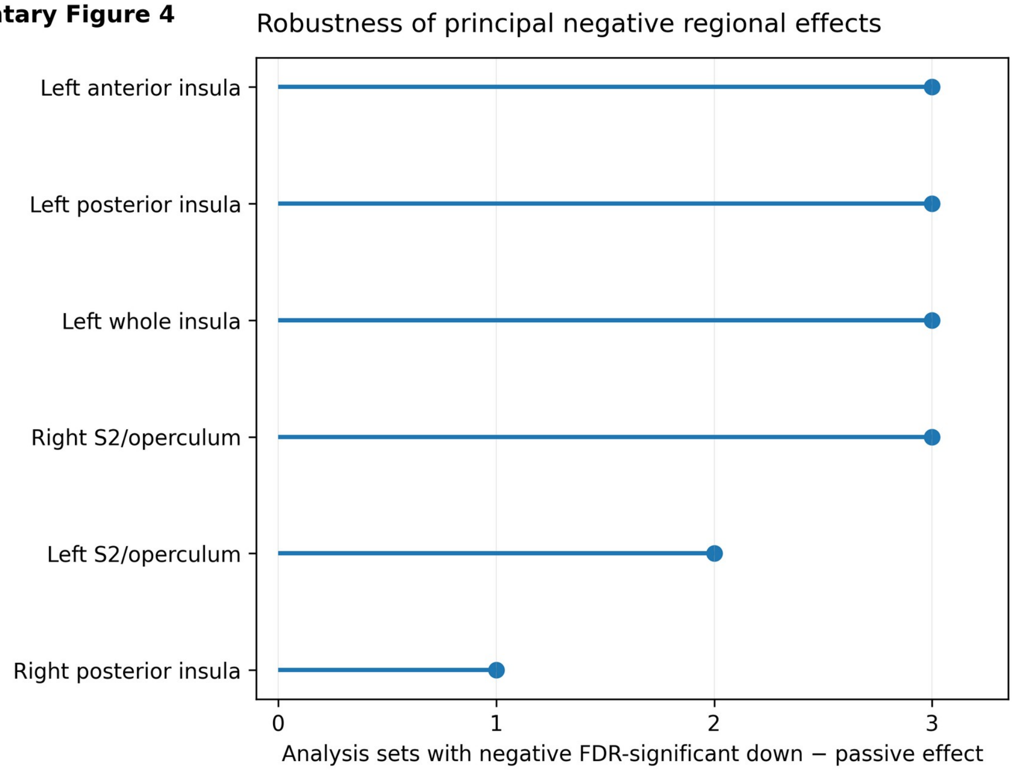


Supplementary Figure 4. Robustness of principal negative regional effects. Points show the number of analysis sets—main, stricter run exclusion and high-motion participant exclusion—in which each down-minus-passive regional estimate was negative and false-discovery-rate significant.

**Supplementary Table S4a. Main and motion-sensitivity down-minus-passive ROI results**

| ROI | Main estimate | Bootstrap 95% CI | p | All-ROI q | Stricter run-exclusion q | High-motion participant-exclusion q |
|---|---|---|---|---|---|---|
| Right S2/operculum | -0.0388 | [−0.0528, −0.0232] | < 0.001 | 0.00064 | 0.002 | 0.009 |
| Left anterior insula | -0.0284 | [−0.0414, −0.0146] | < 0.001 | 0.005 | 0.002 | 0.009 |
| Left whole insula | -0.0225 | [−0.0340, −0.0108] | < 0.001 | 0.008 | 0.005 | 0.009 |
| Left S2/operculum | -0.0276 | [−0.0430, −0.0120] | 0.002 | 0.013 | 0.026 | 0.067 |
| Left posterior insula | -0.0256 | [−0.0411, −0.0102] | 0.003 | 0.019 | 0.005 | 0.033 |
| Right posterior insula | -0.0382 | [−0.0659, −0.0128] | 0.009 | 0.045 | 0.125 | 0.225 |
| Left dlPFC/IFJ | 0.0180 | [0.0052, 0.0301] | 0.010 | 0.045 | 0.064 | 0.225 |

**Supplementary Table S4b. Run-matched sensitivity for selected sensory ROIs**

| ROI | Nearest passive: estimate [95% CI] / p / q / n | Adjacent mean: estimate [95% CI] / p / q / n | Following passive: estimate [95% CI] / p / q / n |
|---|---|---|---|
| Left S2/operculum | −314.7 [−546.2, −83.1] / 0.009 / 0.224 / 32 | −255.8 [−452.8, −58.8] / 0.013 / 0.268 / 33 | −184.1 [−415.0, 46.9] / 0.114 / 0.829 / 32 |
| Right posterior insula | −284.2 [−522.2, −46.2] / 0.021 / 0.224 / 32 | −238.5 [−430.9, −46.1] / 0.017 / 0.268 / 33 | −185.7 [−428.9, 57.5] / 0.130 / 0.829 / 32 |

Note. Estimates are regulation-minus-passive differences in run-mean unscaled FWL-LSS beta coefficients in native fMRIPrep BOLD-intensity units. q values are Benjamini–Hochberg adjusted across all 32 ROI features within each down-regulation-by-comparator family, not across the six displayed cells. Full results for all 256 direction-by-comparator-by-ROI tests are provided in Supplementary Data S1.

**Supplementary Table S5. NPS estimation-path and influential-participant diagnostics**

| Estimation path / quantity | n | Estimate | 95% CI | t | p | FDR q | Diagnostic note |
|---|---|---|---|---|---|---|---|
| Contrast map: down - passive | 33 | −0.00484 | [−0.00697, −0.00278] | −4.45 | < 0.001 | 0.00024 | Stable after excluding one influential participant; path-specific contrast-map units |
| FWL-LSS: up - passive | 33 | -13.844 | [-36.136, 8.448] | -1.27 | 0.215 | 0.307 | No reliable regulation contrast |
| FWL-LSS: down - passive | 33 | -3.345 | [-23.264, 16.575] | -0.34 | 0.735 | 0.735 | No reliable regulation contrast |
| FWL-LSS: mean regulation - passive | 33 | -8.594 | [-25.268, 8.080] | -1.05 | 0.302 | 0.377 | No reliable regulation contrast |
| FWL-LSS: up - down | 33 | -10.499 | [-36.487, 15.488] | -0.82 | 0.417 | 0.463 | No reliable regulation contrast |
| FWL-LSS: passive temperature slope | 33 | 54.317 | [43.405, 65.229] | 10.14 | < 0.001 | < 0.001 | Strong thermal encoding |

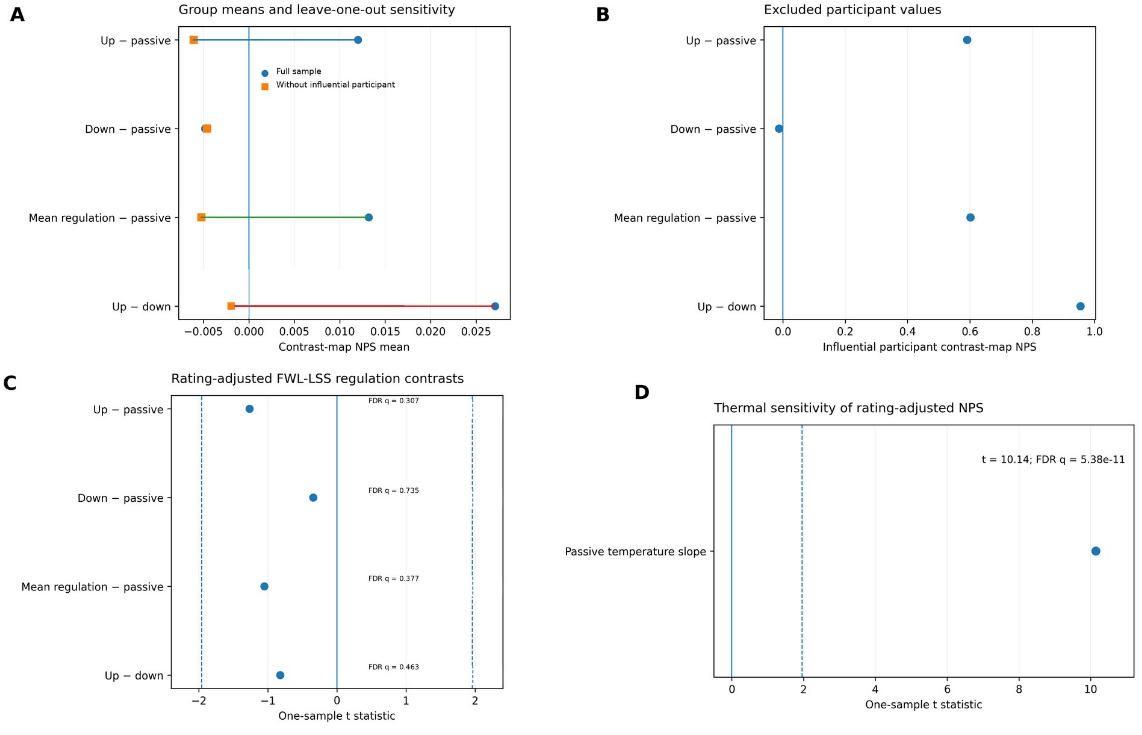


Supplementary Figure 5. NPS estimation-path and corrected trial-wise diagnostics. (A) Contrast-map NPS group means in the full sample and after excluding the influential participant; connected points show the same contrast under the two analysis sets. (B) The corresponding contrast-map NPS values for the excluded influential participant, shown on a separate scale to make leverage visible without compressing the group means. (C) One-sample t statistics for the four primary rating-adjusted FWL-LSS regulation contrasts; dashed lines mark |t| = 1.96 and q values use the reported FDR correction by measure. (D) The passive temperature slope from the same rating-adjusted FWL-LSS analysis, shown separately because its large t statistic would otherwise compress the regulation contrasts. The temperature result confirms strong thermal sensitivity, whereas no regulation contrast was reliable.

### Supplementary Note 4. Exploratory pathway diagnostics and model-based DCM comparison

The exploratory screen comprised four pathway composites and 29 participant-level behavioral variables, yielding 116 post-selection correlation tests for the up-regulation lagged-coupling quantity. Archived materials did not establish a preregistered primary pathway or behavioral endpoint. Accordingly, the selected prefrontal–valuation association is treated as a candidate individual-difference result rather than confirmatory evidence. The complete screening matrix and all 116 correlations are provided in the public reproducibility archive. The transition-level sensitivity analysis is included in repository version 1.1.0 and the associated versioned Zenodo archive (https://doi.org/10.5281/zenodo.23010861).

Matched reverse-direction ($r = 0.265$, $p = 0.136$), synchronous zero-lag ($r = 0.064$, $p = 0.725$), and time-reversed ($r = -0.016$, $p = 0.930$) composites were not associated with success. The forward correlation fell outside both the shared-source within-run permutation and nonzero circular-shift null distributions, but these controls remain conditional on the selected pathway and do not correct the preceding screen. Each participant-direction coefficient was estimated from 10 trials (nine lagged transitions) with five residual degrees of freedom after three predictors and an intercept; split-half reliability was not estimable. Thus, individual-level reliability and out-of-sample reproducibility remain unknown. The negative lagged coefficient is a statistical association between trial-wise beta estimates, not evidence of physiological inhibition, neural signal transmission, or effective connectivity.

Six exploratory five-node DCM variants spanning prefrontal–valuation, affective–descending, and sensory–interoceptive families were re-estimated in SPM25 under MATLAB R2023b. Each retained run was modeled as an independent session. Model-family comparison did not provide convergent evidence for a stable direction-specific mechanism. Configuration files, analysis code, and tabulated parameter/model summaries are included in the reproducibility archive. Large intermediate MATLAB model objects are not redistributed but can be regenerated from the documented inputs and workflows.

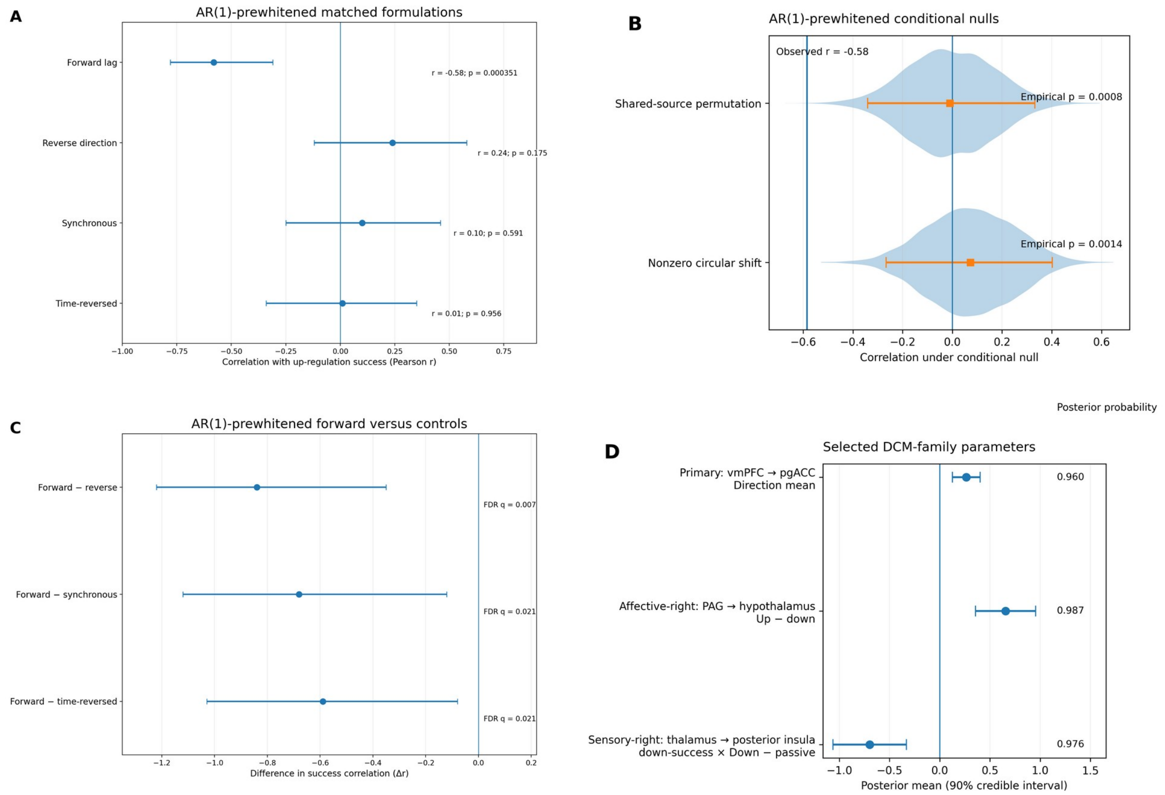


Supplementary Figure 6. AR(1)-prewhitened lagged-coupling sensitivity and exploratory DCM-family results. (A) Correlations between up-regulation success and the AR(1)-prewhitened forward, reverse-direction, synchronous zero-lag and time-reversed composites, with descriptive Fisher-z 95% intervals. (B) Shared-source permutation and nonzero within-run circular-shift null distributions for the AR(1)-prewhitened forward association; the vertical line marks the observed correlation. (C) Participant-bootstrap differences between the AR(1)-prewhitened forward association and matched controls; q values use Benjamini-Hochberg FDR correction across the three comparisons. (D) Selected retained or substantively relevant DCM-family parameters with 90% posterior credible intervals and posterior probabilities. Coupling follow-ups remain conditional on the selected pathway; DCM effects are exploratory model-family sensitivities.

**Supplementary Table S6. Lagged-composite controls and DCM comparison**

| Analysis | Estimate | 95% CI / null interval | p or Pp | Correction or diagnostic |
|---|---|---|---|---|
| Selected exploratory forward composite | r = −0.546 | [−0.764, −0.270] | p = 0.0010 | Archived screening q = 0.0127 |
| Spearman characterization | ρ = −0.549 | [−0.751, −0.253] | p < 0.001 | Post-selection; approximate Fisher-z CI; n = 33 |
| Covariate-adjusted forward model | β = −0.464 | [−0.681, −0.194] | p = 0.0023 | Label-permutation p = 0.0007; all leave-one-out estimates negative |
| Reverse-direction composite | r = 0.265 | [−0.086, 0.558] | p = 0.136 | Matched control; Fisher-z CI; n = 33 |
| Synchronous zero-lag composite | r = 0.064 | [−0.286, 0.398] | p = 0.725 | Matched control; Fisher-z CI; n = 33 |
| Time-reversed composite | r = −0.016 | [−0.357, 0.329] | p = 0.930 | Matched control; Fisher-z CI; n = 33 |
| Shared-source permutation null | Observed r = −0.546 | [−0.341, 0.336] | p = 0.0014 | 5,000 iterations; conditional on selected pathway |
| Nonzero circular-shift null | Observed r = −0.546 | [−0.271, 0.389] | p = 0.0014 | 5,000 iterations; preserved within-run source ordering |
| Forward - reverse | Δr = −0.811 | [−1.178, −0.320] | p = 0.0022 | FDR q = 0.0066 |
| Forward - zero-lag | Δr = −0.609 | [−1.070, −0.011] | p = 0.0470 | FDR q = 0.0470 |
| Forward - time-reversed | Δr = −0.530 | [−0.954, −0.063] | p = 0.0258 | FDR q = 0.0387 |

| Analysis | Estimate | 95% CI / null interval | p or Pp | Correction or diagnostic |
|---|---|---|---|---|
| DCM family: group-average Up - passive | 0 of 6 variants retained | — | — | Reduced full-rank PEB designs |
| DCM family: group-average Down - passive | 0 of 6 variants retained | — | — | Reduced full-rank PEB designs |
| Affective-right DCM: PAG -> hypothalamus, Up - down | Ep = 0.655 | [0.356, 0.954] | Pp = 0.987 | Model-specific exploratory effect |
| Primary DCM: vmPFC/mOFC -> pgACC/rACC, DirectionMean | Ep = 0.264 | [0.126, 0.401] | Pp = 0.960 | Not a regulation-versus-passive contrast |
| Sensory-right DCM: down success x Down - passive, thalamus -> posterior insula | Ep = −0.698 | [−1.062, −0.333] | Pp = 0.976 | Condition-matched exploratory individual-difference effect |

Note. Coupling controls are conditional on the selected pathway and do not correct the screening stage. DCM effects are exploratory; free energies were not compared across regional variants.

### Supplementary Note 5. Finite-cluster, AR(1), and transition-level sensitivity analyses

Finite-cluster behavioral analyses used CR2 standard errors with Satterthwaite degrees of freedom and 9,999 null-imposed Rademacher wild-cluster-bootstrap replications across 33 participant clusters. The standardized up-versus-passive and down-versus-passive instruction effects remained reliable, as did both direction-specific mean-success estimates. The up-minus-down success asymmetry remained null. The passive run-position slope was small and negative, whereas the direction-by-expected-pain interaction was not reliable under either CR2 or wild-cluster inference (Supplementary Table S7a).

**Supplementary Table S7a. Finite-cluster behavioral inference**

| Analysis | Estimate | CR2 SE | Satterthwaite df | CR2 p | Wild-cluster p | Interpretation |
|---|---|---|---|---|---|---|
| Standardized rating: up versus passive | 0.3402 | 0.0766 | 31.99 | < 0.001 | < 0.001 | Instruction-consistent increase retained |
| Standardized rating: down versus passive | -0.2885 | 0.0702 | 32.0 | < 0.001 | < 0.0001 | Instruction-consistent decrease retained |
| Raw rating: up versus passive | 15.3209 | 4.2866 | 31.99 | 0.0011 | < 0.001 | Instruction-consistent increase retained |
| Raw rating: down versus passive | -13.1406 | 3.528 | 32.0 | < 0.001 | < 0.001 | Instruction-consistent decrease retained |
| Mean direction-corrected success | 0.3142 | 0.0551 | 32.0 | < 0.001 | < 0.001 | Positive mean success retained |
| Up-regulation success | 0.3334 | 0.0736 | 32.0 | < 0.001 | < 0.001 | Positive up-regulation success retained |
| Down-regulation success | 0.2949 | 0.0676 | 32.0 | < 0.001 | < 0.001 | Positive down-regulation success retained |
| Up-minus-down success asymmetry | 0.0423 | 0.0885 | 30.88 | 0.6357 | 0.6318 | No reliable direction asymmetry |
| Passive run-position slope | -0.0266 | 0.0116 | 32.0 | 0.0287 | 0.0306 | Small negative passive trend |

| Analysis | Estimate | CR2 SE | Satterthwaite df | CR2 p | Wild-cluster p | Interpretation |
|---|---|---|---|---|---|---|
| Direction-by-expected-pain interaction | -0.1166 | 0.0684 | 31.15 | 0.0983 | 0.0968 | Not reliable under finite-cluster inference |

Note. CR2 p values use Satterthwaite degrees of freedom. Wild-cluster p values use 9,999 null-imposed Rademacher replications. Intercept-only mean tests use the algebraically equivalent participant-cluster sign-flip implementation. Bootstrap confidence intervals were not inverted.

The trial-wise ROI analysis was then repeated with run-specific AR(1) temporal-noise modeling while retaining the same event and nuisance specification. All 66 regulation runs completed successfully, yielding 660 trials and 32 ROI estimates per trial. The AR(1)-prewhitened forward composite was strongly correlated with the primary forward score (r = 0.963) and remained associated with up-regulation success (r = -0.585, p = 0.00035). Reverse-direction, synchronous zero-lag, and time-reversed controls remained nonsignificant. The observed forward association exceeded both conditional null distributions, and all three direct forward-minus-control differences survived FDR correction (Supplementary Table S7b). These analyses remain conditional on the selected pathway and do not repeat the screening family.

A complementary transition-level sensitivity reduced dependence on participant-specific slopes estimated from only nine transitions. Using the archived rating-adjusted AR(1)-prewhitened trial estimates, all 297 up-regulation transitions were modeled directly for vmPFC, mOFC, and the combined vmPFC/mOFC target. The equal-weight three-target success-by-lagged-source interaction was negative in the unadjusted model (beta = -0.121; participant-label permutation p = 0.0136) and became more negative after adjustment for regulation order, passive mean rating, and passive sensory slope (beta = -0.163; label-permutation p = 0.0003). The adjusted effect remained outside a Freedman-Lane conditional-permutation null (p = 0.0102), with participant-cluster bootstrap 95% CI [-0.290, -0.054]. All 33 leave-one-participant-out estimates remained negative (range, -0.223 to -0.134). Target-specific effects were strongest for vmPFC and the combined vmPFC/mOFC definition; mOFC alone was less stable. Random-slope mixed models had the same coefficient direction but produced convergence or boundary warnings for two of three target definitions. This analysis remains conditional on the same-sample selected pathway and is not independent validation.

**Supplementary Table S7b. AR(1)-prewhitened coupling sensitivity**

| Analysis | Estimate | 95% CI or null interval | p | FDR q or diagnostic |
|---|---|---|---|---|
| AR(1) forward composite | r = -0.5848 | [−0.7730, −0.3022] | < 0.001 | n = 33; Fisher-z CI |
| AR(1) reverse-direction composite | r = 0.2419 | [−0.1106, 0.5403] | 0.175 | n = 33; Fisher-z CI |
| AR(1) synchronous zero-lag composite | r = 0.0970 | [−0.2548, 0.4261] | 0.5913 | n = 33; Fisher-z CI |
| AR(1) time-reversed composite | r = 0.0099 | [−0.3345, 0.3520] | 0.9563 | n = 33; Fisher-z CI |
| AR(1) forward score versus primary forward score | r = 0.9631 | [0.9259, 0.9818] | < 0.001 | Score correspondence; n = 33; Fisher-z CI |
| AR(1) shared-source permutation null | Observed r = -0.5848 | [-0.3405, 0.3317] | 0.0008 | 5,000 iterations; conditional on selected pathway |
| AR(1) nonzero circular-shift null | Observed r = -0.5848 | [-0.2658, 0.4022] | 0.0014 | 5,000 iterations; conditional on selected pathway |

| Analysis | Estimate | 95% CI or null interval | p | FDR q or diagnostic |
|---|---|---|---|---|
| AR(1) forward minus reverse | Delta r = -0.8267 | [-1.2206, -0.3239] | 0.0024 | q = 0.0072 |
| AR(1) forward minus synchronous | Delta r = -0.6818 | [-1.1304, -0.1138] | 0.0204 | q = 0.0208 |
| AR(1) forward minus time-reversed | Delta r = -0.5948 | [-1.0192, -0.0919] | 0.0208 | q = 0.0208 |

Note. The shared-source permutation and nonzero circular-shift analyses used 5,000 iterations each. Direct correlation differences used 5,000 participant bootstrap resamples with Benjamini-Hochberg correction across three comparisons. The vmPFC, mOFC, and combined vmPFC/mOFC targets overlap or are nested and form a three-coefficient composite rather than independent anatomical replications.
